# PI3K Signaling Specifies Proximal-Distal Fate by Driving a Developmental Gene Regulatory Network in Sox9+ Lung Progenitors

**DOI:** 10.1101/2021.02.26.433053

**Authors:** Divya Khattar, Sharlene Fernandes, John Snowball, Minzhe Guo, Matthew C. Gillen, Deborah Sinner, William Zacharias, Daniel T. Swarr

**Author notes:** Address correspondence to: Daniel T. Swarr, MD Cincinnati Children’s Hospital Medical Center Division of Neonatology, Perinatal and Pulmonary Biology Department of Pediatrics University of Cincinnati, Cincinnati, OH 45229, USA. These authors contributed equally to this work (Divya Khattar, Sharlene Fernandes).

## Abstract

The tips of the developing respiratory buds are home to important progenitor cells marked by the expression of SOX9 and ID2. Early in embryonic development (prior to E13.5), SOX9+ progenitors are multipotent, generating both airway and alveolar epithelium, but are selective progenitors of alveolar epithelial cells later in development. Transcription factors, including *Sox9, Etv5*, *Irx*, *Mycn*, and *Foxp1/2* interact in complex gene regulatory networks to control proliferation and differentiation of SOX9+ progenitors. Molecular mechanisms by which these transcription factors and other signaling pathways control chromatin state to establish and maintain cell-type identity are not well-defined. Herein, we analyze paired gene expression (RNA-Seq) and chromatin accessibility (ATAC-Seq) data from SOX9+ epithelial progenitor cells (EPCs) during embryonic development. Widespread changes in chromatin accessibility were observed between E11.5 and E16.5, particularly at distal cis-regulatory elements (e.g. enhancers). Gene regulatory network (GRN) inference identified a common SOX9+ progenitor GRN, implicating phosphoinositide 3-kinase (PI3K) signaling in the developmental regulation of SOX9+ progenitor cells. Consistent with this model, conditional ablation of PI3K signaling in the developing lung epithelium *in vivo* and in lung explants *in vitro* resulted in an expansion of the SOX9+ EPC population and impaired epithelial cell differentiation. These data demonstrate that PI3K signaling is required for epithelial patterning during lung organogenesis, and emphasize the combinatorial power of paired RNA-Seq and ATAC-Seq in defining regulatory networks in development.

## Introduction

Morphogenesis of the mammalian lung begins as an outpouching from the embryonic foregut, which branches and extends into the splanchnic mesoderm as the primitive respiratory tubules are formed. The tips of the lung bud are lined by highly proliferative epithelial progenitor cells marked by expression of the genes *Sox9* and *Id2* (Morrisey & Hogan, 2010; Rawlins, Clark, Xue, & Hogan, 2009). Over the course of early lung development, SOX9^+^ progenitor cells give rise to the entire complement of lung epithelium, from proximal airway epithelial cells to distal type I (AT1) and type II alveolar (AT2) epithelial cells (Laresgoiti et al., 2016; Miller et al., 2018; Nichane et al., 2017; Nikolic et al., 2017; Rawlins et al., 2009). Remarkable progress has been made in understanding the signaling pathways (e.g. Wnt, Bmp, Shh, retinoic acid (RA), and FGF signaling) and transcription factors that control development, differentiation, and maturation of the lung epithelium (Morrisey & Hogan, 2010; Swarr & Morrisey, 2015; Whitsett, Kalin, Xu, & Kalinichenko, 2019). Present understanding of how these various signals are integrated to establish respiratory epithelial cell identity remains limited; how this “cellular memory” is disrupted over the course of the human lifespan during homestasis, repair after injury, and in various lung diseases is an area of active investigation.

Chromatin state serves as a medium through which various developmental and external stimuli are integrated into comprehensive cellular programs which enable regulatory proteins access to activate or repress gene expression. Epigenetic programs controlling progenitor cell identies are maintained in spite of repeated rounds of cellular division, fluctuating transcription factor levels, or dynamic inputs from various signaling pathways (Klemm, Shipony, & Greenleaf, 2019; Sheikh & Akhtar, 2019). Chromatin accessibility changes dynamically throughout development, to establish and maintain cellular identity in a variety of cell types and organ systems (Gorkin et al., 2020; Trevino et al., 2020). The extent to which changes in chromatin accessibility contribute to lung development as a diversity of mature epithelial cells differentiate from embryonic progenitors remains incompletely understood.

Chromatin accessibility, or “the degree to which nuclear macromolecules are able to physically interact with chromatinized DNA” can now be measured in a relatively small number of cells using Assay for Transposible-Accessible Chromatin using sequencing (ATAC-Seq) (Buenrostro, Giresi, Zaba, Chang, & Greenleaf, 2013; Klemm et al., 2019), wherein Tn5 transposase inserts Illumina sequencing adapters into accessible regions of chromatin. Although accessible chromatin regions account for a minority of the genome (2-3%), they account for a high proportion of the genomic loci bound by transcription factors, and are highly correlated with cis-regulatory elements (CREs) identified by other methods, such as promoters, enhancers, insulators, and silencers (Buenrostro et al., 2013; Buenrostro, Wu, Litzenburger, et al., 2015).

Herein, we used paired RNA-Seq and ATAC-Seq to define the chromatin accessibility landscape of the developing SOX9+ lung epithelial progenitor cells at multiple developmental timepoints. Acessible chromatin regions in SOX9+ cells were highly correlated with lung-specific histone post-translational modifications (PTMs) found on validated cis-regulatory elements. Dynamic changes in these accessible chromatin regions were correlated with gene expression changes observed during the course of SOX9+ lung epithelial cell differeniation. Chromatin accessibility changes were observed in or near genes associated with phophatidylinositol 3-kinase (PI3K) signaling. Inhibition or genetic ablation of PI3K signaling caused expansion of the SOX9+ progenitor cell population at the expense of mature lung epithelial cell differentiation. Our results demonstrate the power of epigenomic analysis to identify new roles for gene regulatory networks and signaling pathways in lung epithelial development, highlighting the previously unappreciated role of PI3K signaling in SOX9+ cell differentiation and proximal-distal cell fate specification.

## Results

### The multi-potent SOX9+ lung epithelial progenitor cell (EPC) population

Shortly after lung specification occurs, highly proliferative distal progenitor cells marked by the expression of SOX9+ appear at the tips of the growing lung buds (Figure 1A-I). A suite of transcription factors (eg, Id2, Etv5, Foxp1/2, Mycn) play critical roles in the development of the SOX9+ distal tip progenitor cell population (Figure 1D) (Morrisey & Hogan, 2010; Swarr & Morrisey, 2015). Several lines of evidence, including lineage tracing and 3D culture model systems, demonstrate that early in development SOX9+ progenitor cells are multi-potent, generating both airway and distal alveolar epithelium (Laresgoiti et al., 2016; Nichane et al., 2017; Nikolic et al., 2017; Rawlins et al., 2009). In order to orthogonally validate these observations and gain insight into the transcriptional changes occurring during these developmental decisions, we peformed pseudotime analysis (Monocle) of RNA-Seq data from mouse lung epithelium from E11.5 to E16.5 (Trapnell et al., 2014). Starting from E11.5, two distinct pseudotime branches arise from SOX9+ progenitors, terminating in airway and distal lung epithelial cells at E16.5 (Figure 1J-K, Supplemental Figure 1A-B). We used Monocle to find modules consisting of co-regulated genes during distal differentiation, revealing 26 unique gene modules. A subset of these gene modules overlap with pseudotime analysis and correspond with specific aspects of SOX9+ epithelial differentiation (Figure 1L-M, and Supplemental Figure 1). One such gene module consists of genes highly expressed in E11.5 lung and are preserved the distal differential branch of the Monocle pseudotime analysis; these genes are involved in lung epithelial development and early branching including *Sox9*, *Wnt7b*, *Bm4* and *Hmga2*, validating the significance of both the pseudotime analyses and gene module predictions while also identifying other coordinately expressed genes that may be involved in EPC differentiation (module 8, Figure 1L-M). Other modules highlight genes that likely regulate the proliferating progenitor cells on the same central branch to distal epithelium (module 16, Supplemental figure 1E,G), and genes involved in the functions of differentiation of airway epitheilum (modules 4 &12, Supplemental Figure C-D,F). These data are consistent with gene set enrichment analysis of bulk RNA-Seq data at E11.5 and E16.5 (Supplemental Figure 2 & Supplemental Table 1).

**Figure 1.**
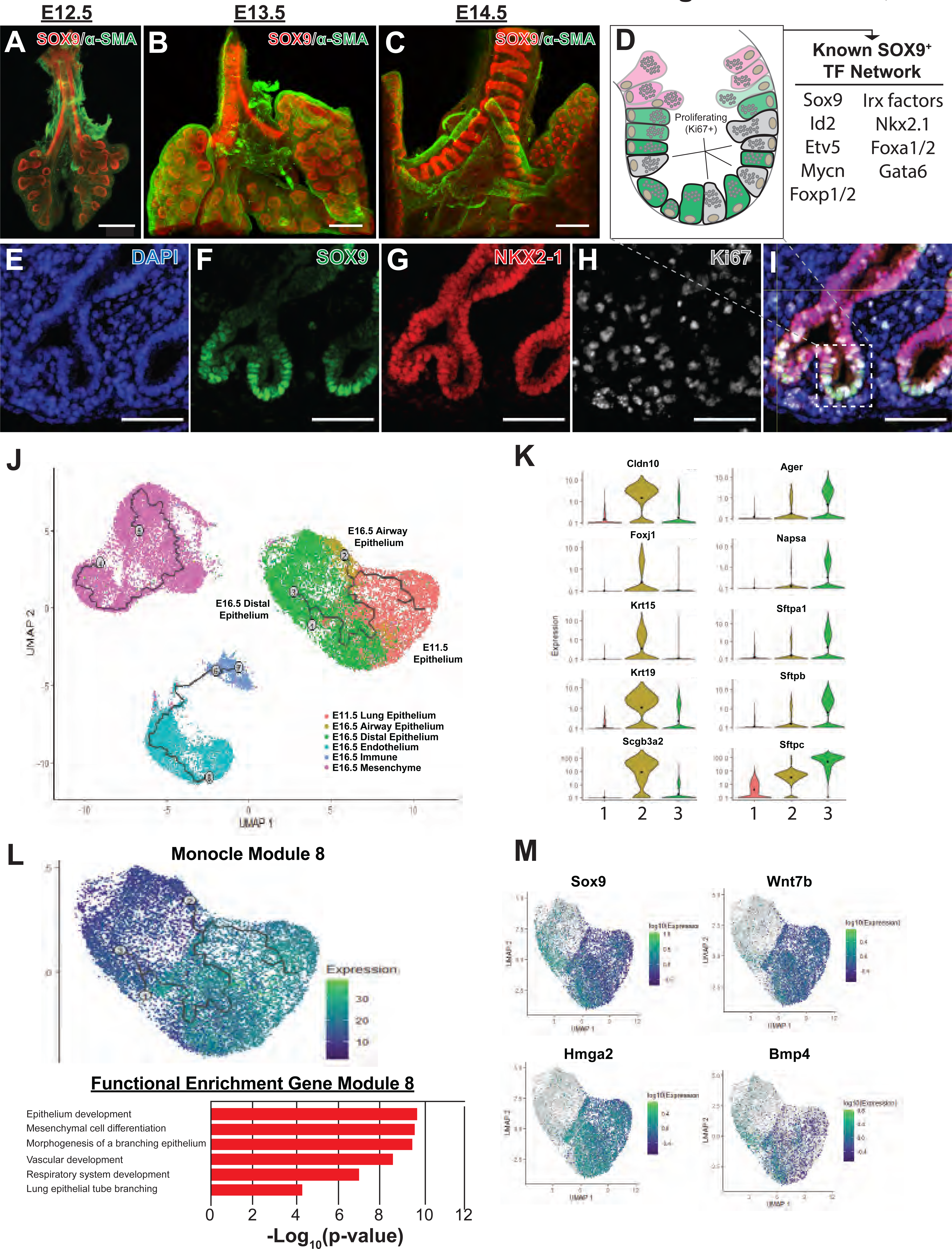
The SOX9+ multipotent lung epithelial progenitor cell population. (A-C) Whole mount confocal imaging of embryonic lungs isolated from SOX9-GFP reporter mice at E12.5 (A), E13.5 (B), and E14.5 (C) show SOX9 expression at the distal branch tips and in the tracheal mesenchyme. (D) Schematic summarizing the transcription factors known to play an important role in the development and differentiation of SOX9+ progenitors. (E-I) Immunofluoresence microscopy shows high levels of SOX9 staining in the epithelium of the distal lung tips, with a high percentage of Ki67+ proliferating cells early in development. (J-M) Monocle3 lineage analyses was performed on E11.5 lung epithelial cells and E16.5 lung cells. (J) Lineage analyses predict E11.5 lung epithelial cells give rise to both proximal and distal lung epithelial cells at E16.5. (K) Known distal and proximal genes markers are expressed in their corresponding cell clusters at E16.5 but not E11.5, validating cluster identification. Cluster labels = 1 – E11.5 lung epithelium; 2 – E16.5 airway epithelium; 3 – E16.5 distal epithelium. (L) Monocle 3 analysis of isolated epithelial populations identifies multiple gene modules with specific expression patterns. One module of gene expression is consistent with the predicted distal differentiation trajectory and consists of genes related to process important to lung development, including epithelial/respiratory system branching and development. (M) The expression patterns of example genes in this module are visualized and include genes known to be essential for proper distal lung development including Sox9, Wnt7b, Hmga2 and Bmp4. Scale bars: 200 μm (A-C), 50 μm (E-I).

In order to evaluate the landscape and dynamics of cis-regulatory DNA elements associated with these transcriptional changes occurring during development of EPCs, SOX9+ progenitor cells (SOX9-GFP^+^, CD326^+^, CD31^-^, CD45^-^, 7-AAD^-^) were isolated by fluorescence-activated cell sorting (FACS) from a transgenic reporter mouse (containing an EGFP allele driven by the Sox9 promoter element) (Gong et al., 2003) (Supplemental Figure 3). SOX9+ progenitors were then subjected to bulk RNA-Seq and ATAC-Seq. E11.5 and E16.5 were chosen to capture cells prior to, and shortly after, the period of proposed restriction from multi-potent progenitor to distal alveolar progenitor cells (Frank et al., 2019; Nichane et al., 2017; Nikolic et al., 2017; Rawlins et al., 2009) (Figure 2A).

**Figure 2.**
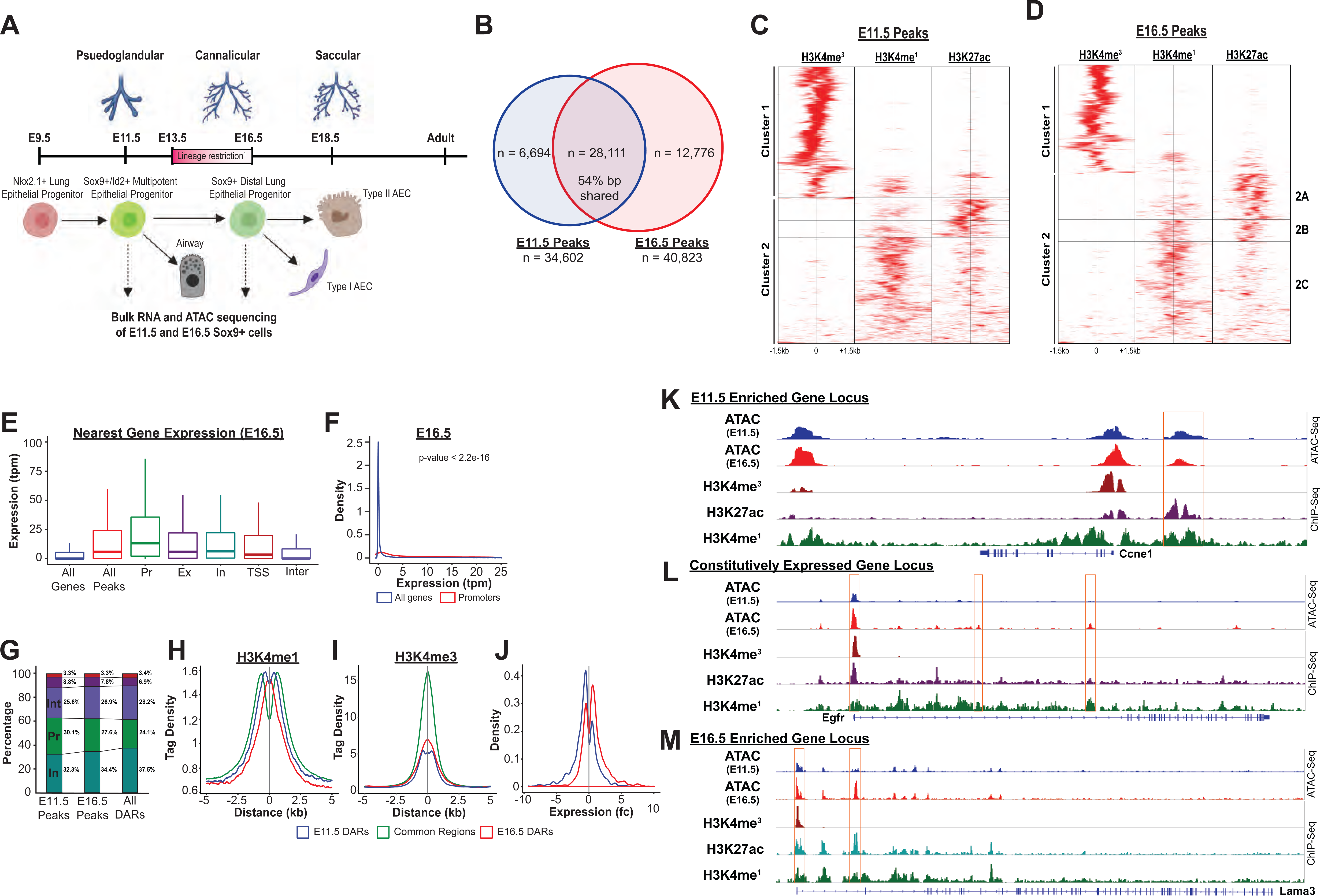
The SOX9+ progenitor cell chromatin accessibility landscape. (A) Schematic strategy for isolation of SOX9+ EPC. Cells were sorted at E11.5 and E16.5 and taken for RNA-seq and ATAC-seq analysis, which correspond to pseudoglandular and late saccular stages of lung development. Notable timepoints of lineage restriction of the SOX9+ EPC population and notable differentiated cells arising from EPCs are noted. (B) More than 34,000 regions of open chromatin were identified by ATAC-Seq at each developmental timepoint. (C-D) ChIP-Seq tag densities for H3K4me^3^, H3K4me^1^, and H3K27ac were plotted and clustered using Homer and R, respsectively. H3K4me^3^ tag density corresponds to the accessible promoter regions, and H3K4me^1^ with or without H3K27ac correspond to active and “decommissioned” enhancer regions, respectively. (E-F) Gene expression correlates to nearby open chromatin, especially in the promoter region. Pr – Promoter; Ex – Exonic; In – Intronic; TSS – Transcriptional Stop Site; Inter – Intergenic (G) Differentially accessible chromatin regions (DARs) were more likely to be located with intronic (In) and intergenic (Int) regions instead of promoters. (H) H3K4me^1^ peak enrichment is found at the center of DARs. (I) Common regions were more enriched for H3K4me^3^ signals compared to DARs as either timepoint. (J) E11.5 and E16.5 DARs correlate with expression of nearby genes by RNA-seq. (K-L) Examples of three categories of genomic loci are shown - (K) Decreasing levels of accessibility over time for progenitor genes, (L) Changing accessibility without change in gene expression for housekeeping genes, and (M) Increasing accessibility at later developmental timepoints for distal genes.

**Figure 3.**
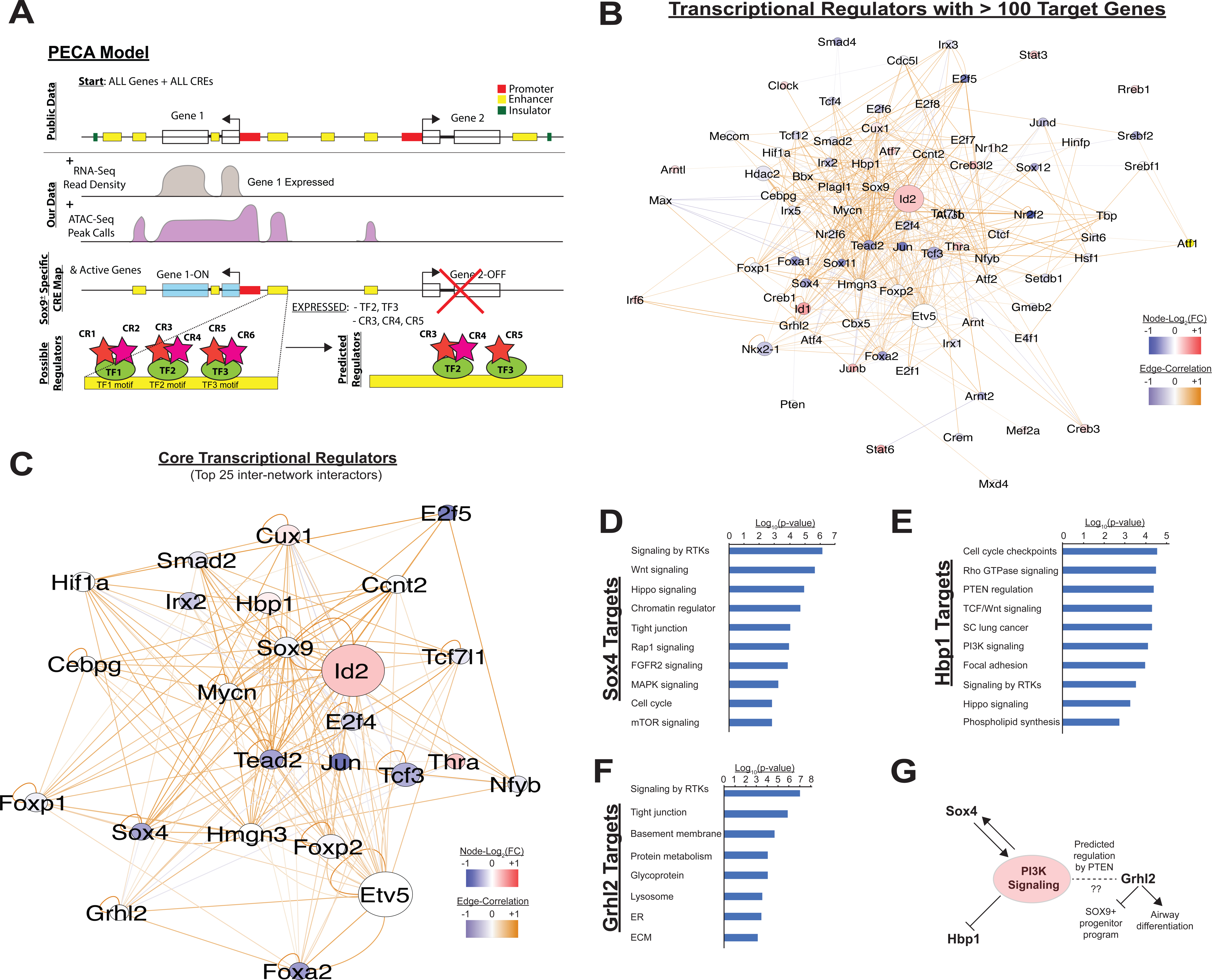
Paired Expression and Chromatin Accessibility (PECA) modeling of SOX9+ progenitor cells. (A) The computational model paired expression and chromatin accessibility (PECA) was used to develop a regulatory model based on these paired RNA-Seq and ATAC-Seq datasets. Potential active cis-regulatory elements, key transcription factor (TF) networks, and chromain regulatory (CR) active in the SOX9+ lung epithelial progenitor cells are predicted. (B) The co-regulatory relationships between transcription factors with more than 100 predicted target genes are shown. Network nodes (TFs) are color coded according to their expression log2 fold-change between E11.5 to E16.5 (blue – decreasing expression, red – increasing expression), and network edges are color coded by the correlation coefficient between the two indicated TFs (blue and organge indicate negative & positive regulatory relationships, respectively). (C) The transcription factors with the top 25 highest degree of network interactions in EPCs. (D-E) The TFs *Hbp1* and *Sox4* interact with PI3K signaling. Gene ontology analysis of predicted targets of these TFs in the SOX9+ progenitor cell network are shown. (F) *Grhl2* has previously been shown to promote airway differentiation, and its loss leads to expansion of the SOX9+ progenitor population. Our network data predict regulation of *Grhl2* expression through the PI3K signaling pathway. (G) Putative relationship between the PI3K signaling pathway and several key nodes of the SOX9+ progenitor cell gene regulatory network.

### Accessible regions of chromatin in SOX9+ EPCs are active cis-regulatory elements that correlate with neighboring gene expression

Analysis of chromatin accessibility by ATAC-Seq identified over 30,000 regions of open chromatin at each timepoint (E11.5, n = 34,602; E16.5, n = 40,823)(Figure 2B); only peaks replicated in independent biological samples were included for further analysis (Supplemental Figure 4A-B, Supplemental Tables 2-3). The genome-wide distribution of ATAC-Seq peaks was consistent with previously published chromatin accessibility studies, showing a high-degree of enrichment near gene transcriptional start sites (TSS) (Supplemental Figure 4C-H). The presence of ATAC-Seq peaks positively correlated with expression the nearest gene in the RNA-Seq, and this correlation was strongest for peaks within promoter regions. Moreover, peaks only detected at a single developmental timepoint were associated with higher levels of expression of the nearst gene at that same developmental stage (for example, genes nearest to peaks unique to the E11.5 developmental stage overall were observed to have higher expression levels at E11.5 compared to E16.5). (Figure 2E-F, Supplemental Figure 5A-E)

To determine whether these accessible regions of chromatin had shared features of active cis-regulatory DNA elements (CREs), we examined H3K4me^3^, H3K4me^1^, and H3K27ac ENCODE ChIP-Seq read density (embryonic mouse, whole lung) across all identified ATAC-Seq peaks (Consortium, 2012; Davis et al., 2018). Significant enrichment was observed for H3K4me^3^ read density across peaks predicted to reside within gene promoter regions. (Figure 2C-D and Supplemental Figure 5F-G) A second subset of ATAC-Seq peaks were enriched for either both H3K4me^1^ and H3K27ac read density, or H3K4me^1^ ready density alone, consistent with “active” or “decommissioned” enhancer states, respectively (Jadhav et al., 2019). (Figure 2C-D and Supplemental Figure 5F-G) Taken together, these data suggest that regions identified by ATAC-Seq represent a bona fide map of active cis-regulatory elements (e.g. promoters, active and decommissioned enhancers, etc) within the SOX9+ lung epithelial progenitor cells, and that these regulatory elements are regulating the transcriptional dynamics observed during EPC development.

### Developmental changes in SOX9+ EPC chromatin accessibility preferentially occur outside of promoter regions

Nearly a quarter of accessible chromatin regions were unique to each developmental timepoint, a remarkable observation given that the two cell populations are SOX9+ lung epithelial progenitor cells separated by just 5 days of development (Figure 2A). We used the MACS2 bdgdiff tool to identify regions of chromatin that were more accessible at E11.5 (n = 18,582), more accessible at E16.5 (n = 11,980), or did not undergo significant changes in accessibility between the two timepoints (n = 35,312) (hereafter referred to as E11.5-differentially accessible regions (DARs), E16.5-DARs, and common regions, respectively) (Supplemental Figures 4-6). As was observed for all peaks, both sets of DARs and common regions were significantly enriched within 2kb of transcriptional start sites (Supplemental Figure 5H-I). However, compared to peaks that did not undergo changes in chromatin accessibility, DARs were significantly more likely to be found within intronic and intergenic regions, and were less likely to be found within promoters (Figure 2G and Supplemental Figure 5I). DARs at both E11.5 and E16.5 exhibited peak H3K4me^1^ read density in close proximity to the center of the DAR. In contrast, there was relative de-enrichment of H3K4me^1^ read density at the center of common regions (and peaks flanking the common region by ∼500bp on each side) (Figure 2H). Similarly, read density for the promoter mark H3K4me^3^ was significantly enriched and centered on common region peaks, compared to DAR peaks (Figure 2I). At the genome-wide level, changes in chromatin accessibility were significantly correlated with differential gene expression. Genes adjacent to regions of chromatin more accessible at E11.5 (E11.5-DARs) were significantly more likely to have higher mRNA expression at E11.5 compared to E16.5, and vice versa (Figure 2J). Examples of differentially accessible chromatin regions flanking or within two highly differentially expressed genes (e.g. Ccne1, Lama3), and common accessible regions within a stably expressed gene locus (e.g. Egfr), are shown in Figure 2K-M. These data support the concept that regions of chromatin accessibility identified in this study contain active CREs that influence neighboring gene expression. Moreover, although chromatin remodeling occurs across all types of cis-regulatory elements during development of SOX9+ lung epithelial progenitors, developmental changes in chromatin accessibility were preferentially located outside of the promoter regions at distal cis-regulatory elements (e.g. enhancers). Collectively, these data also emphasize that chromatin accessibility at distal CREs such as enchancers likely plays a key and incompletely explored role in directing lung epithelial development.

**Figure 4.**
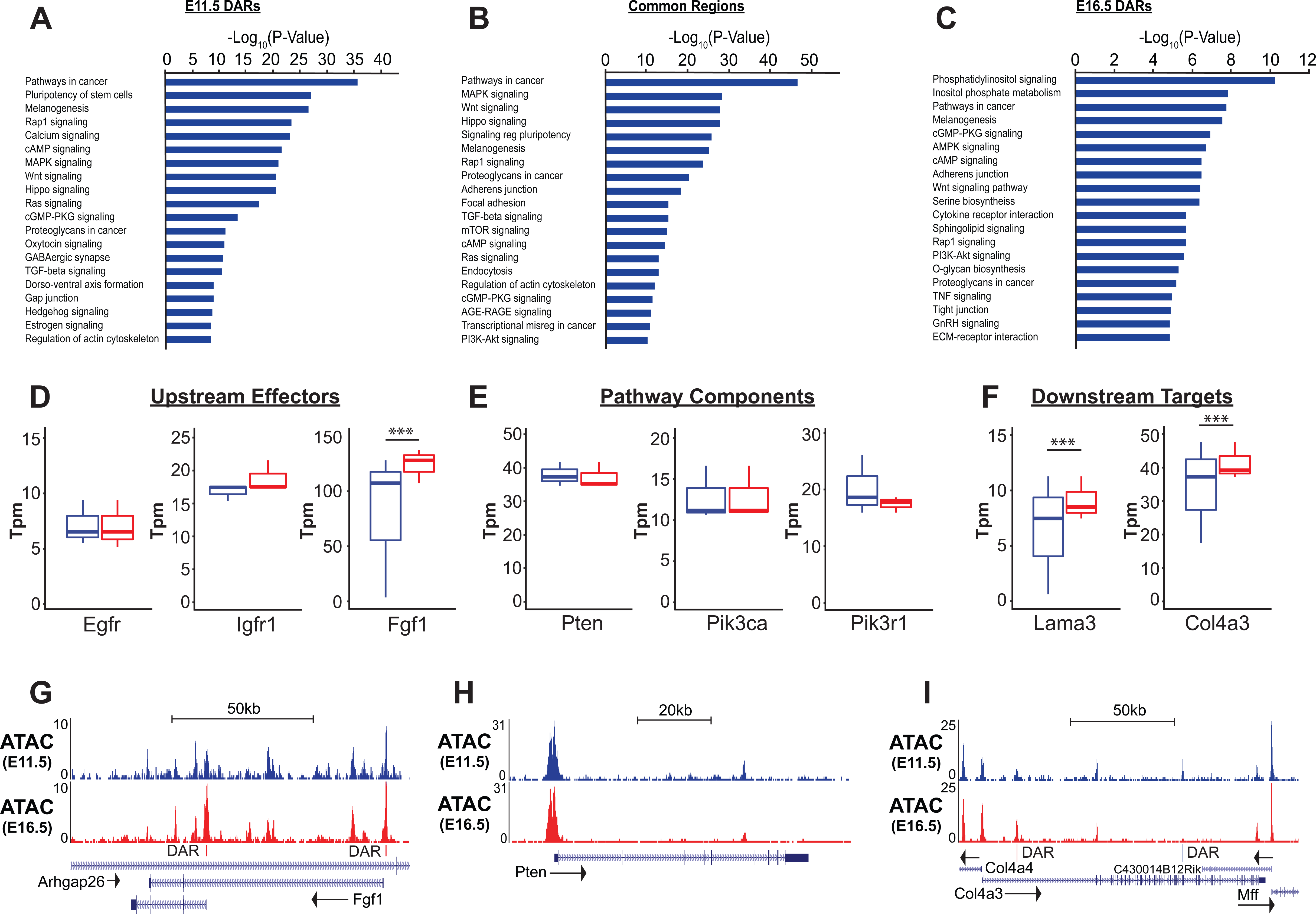
Differentially accessible chromatin regions associated with PI3K-pathway related genes. (A-C) Gene ontology analysis for genes nearest E11.5 & E16.5 differentially accessible regions (DARs) and common regions are shown. Pathways regulating lung epithelial development, such as Wnt, Hippo, and Hedgehog signaling are identified. A number of GO catatories associated with PI3K signaling were over-represented in genes adjacent to E16.5 DARs, as well as those nearby common regions. (D-I) Examples include proteins upstream of PI3K signaling (D,G), core pathway components (E,H), and known downstream targets of PI3K sigaling (F,I). Many of the downstream targets are components of epithelial basement membrane, which increase as development proceeds. (G-I) Examples of DARs correlated with gene expression changes for *Fgfr1* and *Col4a3* are shown. *Pten*, which does not change in expression between E11.5 and E16.5 has common accessible regions of chromatin (in promoter and 5^th^ intron). *** p < 0.001.

**Figure 5.**
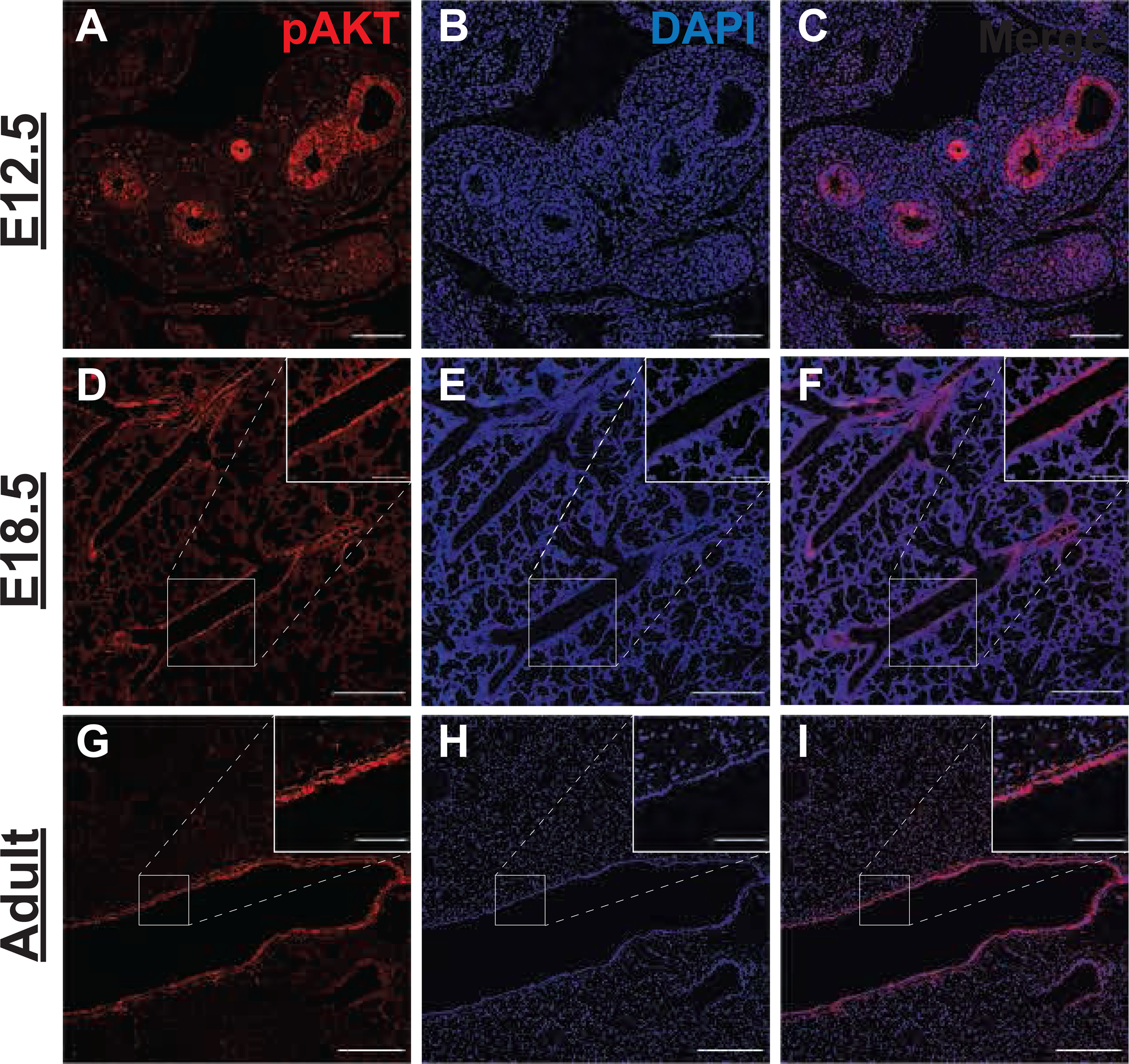
Phospho-AKT staining in airway epithelial cells. (A-C) Strong expression of phospho-AKT at Ser473 (pAKT) was observed in the early lung epithelium at E12.5, with much lower levels in the mesenchyme. (D-F) By E18.5, highest levels of pAKT were detected in epithelial cells lining the large conducting airways. (G-I) Strong pAKT staining was observed in the conducting airway epithelium, with notable expression in the sub-epithelial airway mesenchyme (see inset). Scale bars: 100 μm (A-C), 250 μm (D-I), 100 (inset D-F), 50 μm (inset G-I).

**Figure 6.**
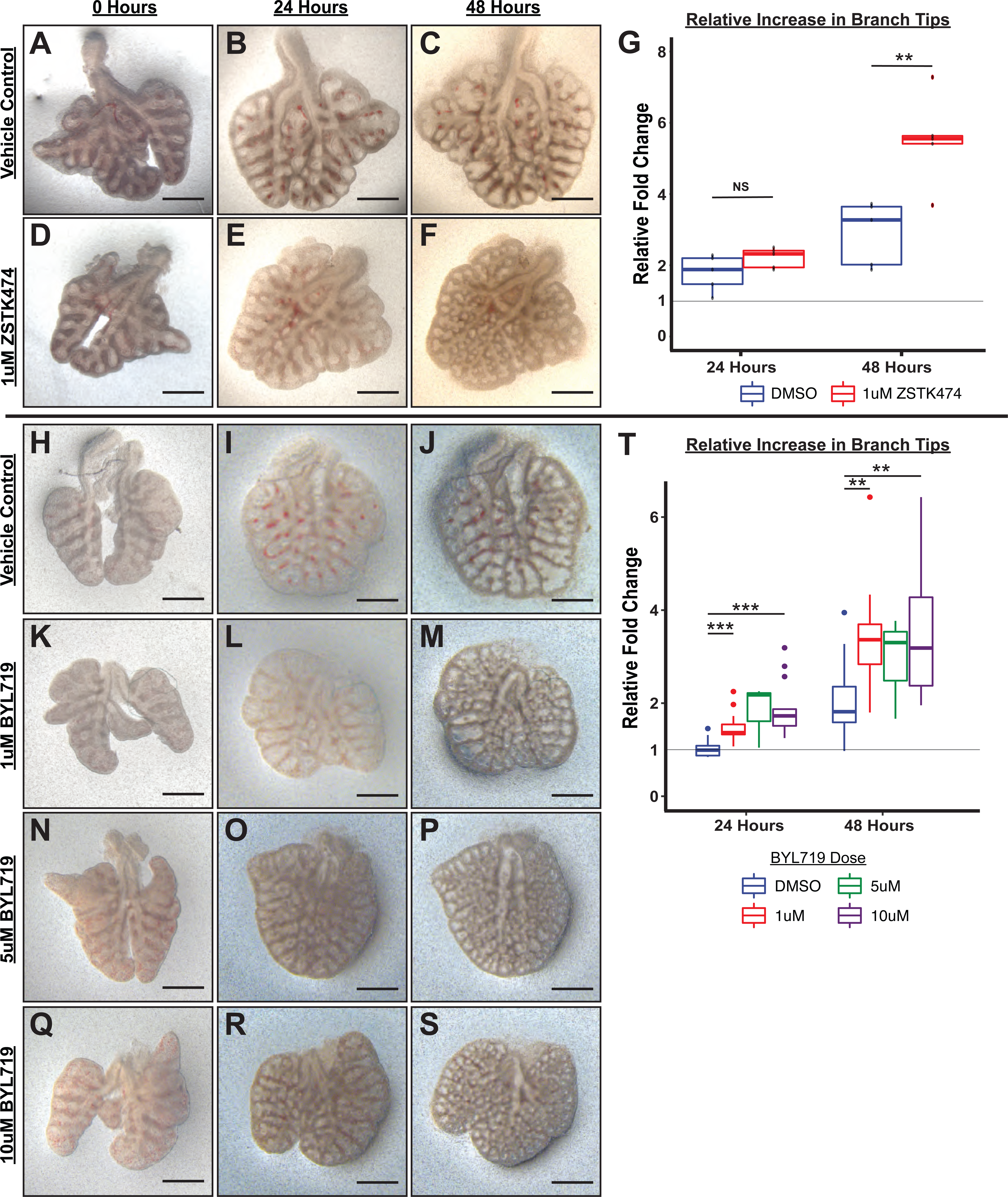
Inhbition of PI3K signaling increases the number of distal lung tips. Lungs were isolated from wild-type mouse embryos at E12.5 and were cultured for 48 hours in the presence of one of two pan-class I PI3K pathway inhibitors or vehichle control. Following treatment with either ZSTK474 (A-G), or BYL719 (A-G), the number of lung branch tips significantly increases. Scale bars: 500 μm. ** p < 0.01, *** p < 0.001.

### Chromatin remodeling is enriched near loci associated with PI3K signaling

Although analysis of gene expression alone enables predictions regarding signaling pathways and transcription factor networks driving transcriptional changes, we hypothesized that adding an understanding the cis-regulatory landscape within developing SOX9+ lung epithelial cells would provide additional mechanistic insights into how these trans-activating factors promote and maintain cell differentiation (Klemm et al., 2019; Trevino et al., 2020). To this end, we used the Paired Expression and Chromatin Accessibility (PECA) computational model to build a gene regulatory network for the SOX9+ EPC population at E11.5 and E16.5 (Duren, Chen, Jiang, Wang, & Wong, 2017). At the epigenomic level, this model compares open regions of chromatin to a map of known cis-regulatory elements identified through large-scale epigenomics consortia (e.g. ENCODE), TF motifs detected within these regions, and TF expression data derived from expression data to eliminate spurious motif elements (for example, motifs for transcription factors that are not expressed at the mRNA level in the paired mRNA-seq dataset). The model then compares these open elements to paired mRNA-seq data in order to make predictions about active TFs, the genes they regulate, the cis-regulatory elements that they bind to, and the chromatin regulatory complexes with which they interact (Figure 3A). The interactions between TFs predicted to regulate more than 100 target genes are shown in Figure 3B. Transcription factors with the highest degree of interconnectivity (top 25 TFs) are shown in Figure 3C. Well-studied transcription factors, including those highlighted in Figure 1D, such as *Id2*, *Irx* and *Foxp factors*, *Etv5*, and *Foxa2* appear as central nodes.

Additional TFs play important roles in transcription generally (e.g. *E2F*, *Jun*). However, multiple other TFs that appear in this central network have been less-well studied in the contex of lung epithelial development. Intriguingly, several of these key nodes have previously shown to interface with the PI3K signaling pathway (Figure 3D-G). The transcription factor *Sox4*, which has been implicated in the pathogenesis of non-small cell lung cancer (NSCLC) and was recently found to be enriched in a regeneration associated pre-alveolar type-1 cell transitional state (PATS), has been shown to both regulate and have its own expression regulated by PI3K signaling (Bilir et al., 2016; Kobayashi et al., 2020; Mehta et al., 2017; Ramezani-Rad et al., 2013; Wang, Hao, Pan, Qian, & Zhou, 2015). Another central node in this network, *Hbp1,* is a known repressor of beta-catenin transactivation, but is also negatively regulated by PI3K signaling (Bollaert et al., 2018; Coomans de Brachene et al., 2014). Finally, *Grhl2*, a third central node in this network, was predicted to be regulated via the PI3K signaling component *Pten*. Previous work has shown *Ghrl2* promotes airway differentiation in the lung, and that loss of *Grhl2* in the lung leads to expansion of the SOX9+ EPC population (Kersbergen et al., 2018).

To complement the gene-regulatory network model predicted by PECA, we performed gene set enrichment analysis (GSEA) for both RNA-Seq and ATAC-Seq data sets, using the nearest neighbor gene-peak pair for ATAC data. KEGG pathway analysis of RNA-Seq data identified biological processes and pathways consistent with known regulators of lung epithelial development. At E11.5, gene categories involving cell proliferation (e.g. “cell cycle”, “homologous recombination”), protein synthesis (e.g. “ribosome”, “biogenesis”), and “signaling pathways regulating pluripotency” are most enriched. By E16.5, processes related to develop of functions associated with the mature distal lung epithelium, such as surfactant production (e.g. “phagosome”, “lysosome”, “sphingolipid metabolism”, “protein export), epithelial barrier function (e.g. “ECM-receptor interaction”, “focal adhesion”, “collective duct secretion”), and a switch in energy metabolism (e.g. “oxidative phosphorylation”, “citrate/TCA cycle”) are apparent (Supplemental Figure 2).

While similar biological processes were enriched in both the ATAC-Seq and RNA-Seq data, analysis of ATAC-Seq data identified activation of a number of signaling pathways that were less apparent by RNA analysis alone. Signaling pathways known to be important in lung epithelial development, such as Wnt, Hedgehog, Notch, and Hippo signaling were identified in both analyses. However, analysis ATAC-Seq data identified a potential role for a number of additional pathways in the development and differentiation of the SOX9+ lung epithelial progenitor cells, including cAMP, *Rap1*, *Ras*, MAPK, TGF-beta, and, importantly, PI3K-mTOR signaling (Figure 4A-C and Supplemental Figure 2B-C).

Consistent with both the PECA and GSEA analyses, genes nearest to increasingly accessible regions of chromatin (E16.5-DARs) as EPC maturation proceeded were most strongly enriched in genes associated with phosphatidylinositol (KEGG mmu04070) and PI3K signaling (KEGG mmu04151). Genes nearest to common accessible regions were also significantly enriched in these same pathways (Figure 4B). Components of the PI3K signaling pathway (eg. *Pik3ca*, *Pikap1*, *Pik3r1*, *Pten*), upstream ligand/receptors (eg. *Egfr*, *Fgf1*, *Igf1r*) and known downstream targets of PI3K signaling were associated with common accessible regions and E16.5 DARs (Figure 4D-F). While expression of many of core pathway components did not change between E11.5 and E16.5, expression of a number of potential upstream receptor/ligands (e.g. *Igfr*, *Fgf1*) and downstream targets of PI3K signaling increased, including major components of the epithelial basement membrane (e.g. *Lama3*, *Col4a3*). For example, differentially accessible regions within two promoter regions of the *Fgf1* locus and a E16.5-DAR within the first intron of the *Col4a3* locus are shown in Figure 4G,I. In contrast, common accessible chromatin regions are seen within the promoter and 5^th^ intron of *Pten*, which is stably expressed between E11.5 and E16.5 in the SOX9+ lung epithelium (Figure 4H). Taken together, these data suggested that the dynamic changes in chromatin state observed during SOX9+ EPC development were modulated by PI3K signaling, an observation only possible with combined chromatin and RNA analysis.

### Inhibition of PI3K Signaling in lung explants causes increased branching and expansion of the SOX9+ EPC population

In order to evaluate the hypothesis that PI3K pathway activation occurs differentially throughout the developing lung epithelial cell population, we measured PI3K pathway activity by detecting AKT protein phosphorylated at Ser-473 (pAKT) in the embryonic mouse lung (Alessi et al., 1996; S. Zhang et al., 2017). The highest levels of pAKT expression, and PI3K pathway activation, were observed in the proximal lung epithelium as early as E12.5 and persisted in the airway epithelium through adulthood. Much lower levels of pAKT expression and PI3K activation were observed in the distal lung epithelium and mesenchyme. These observations are consistent with a recent published study of early postnatal lung, which demonstrated highest levels of pAKT at postnatal day 5 within the airway epithelium (K. Zhang et al., 2020). These data support the concept that a proximal-to-distal gradient of PI3K signaling within the lung epithelium may be required for lung epithelial differentiation and proper establishment of epithelial cell-type identity.

To test the hypothesis that PI3K signaling influences growth and differentiation of the lung epithelium, we began by using an *in vitro* lung explant model to quantify branching and cellular differentiation in the presence of two independent pan-class I PI3K inhibitors (ZSTK474, BYL719). Lungs were harvested at E12.5 by microdissection and cultured for 48 hours in Transwell inserts (Corning) in the presence of either inhibitor or vehicle control. We observed a significant increase in lung branching after 48 hours in culture with PI3K inhibition (Figure 6). Concordantly, inhibition of PI3K signaling increased the proportion of SOX9+/NKX2.1+ lung epithelial cells and increased *Sox9* mRNA measured by qPCR (Figure 7A-J). Despite increased numbers of branch tips, *Sox9* mRNA, and numbers of SOX9+ lung epithelial cells, a decreased number of proliferating (Ki67+) cells was observed in both the epithelium and mesenchyme (Figure 7K-P). Immunofluorescence staining of mature surfactant protein C (SFTPC), and qPCR for *Sftpc* and *Lamp3* mRNA, both normally highly expressed in the mature AT2 cells, were significantly decreased following PI3K pathway inhibition (Figure 7V-CC). Athough *Scgb3a2* mRNA was significantly decreased following PI3K inhibition, no significant changes in the expression of *Sox2* mRNA was observed, and the early developmental stage did not permit interrogation of expression changes in more mature airway epithelial cell markers (Figure 7FF-GG). Taken together, these data suggested that inhibition of PI3K signaling early in lung development causes expansion of the SOX9+ lung epithelial progenitor cell population at the expense of distal alveolar epithelial maturation. Since our combined ATAC and RNA data suggested that PI3K signaling in the epithelium was the crucial modulator of these effects, we turned to an *in vivo* genetic model to directly assess the epithelial-specific requirement for PI3K signaling during lung development.

**Figure 7.**
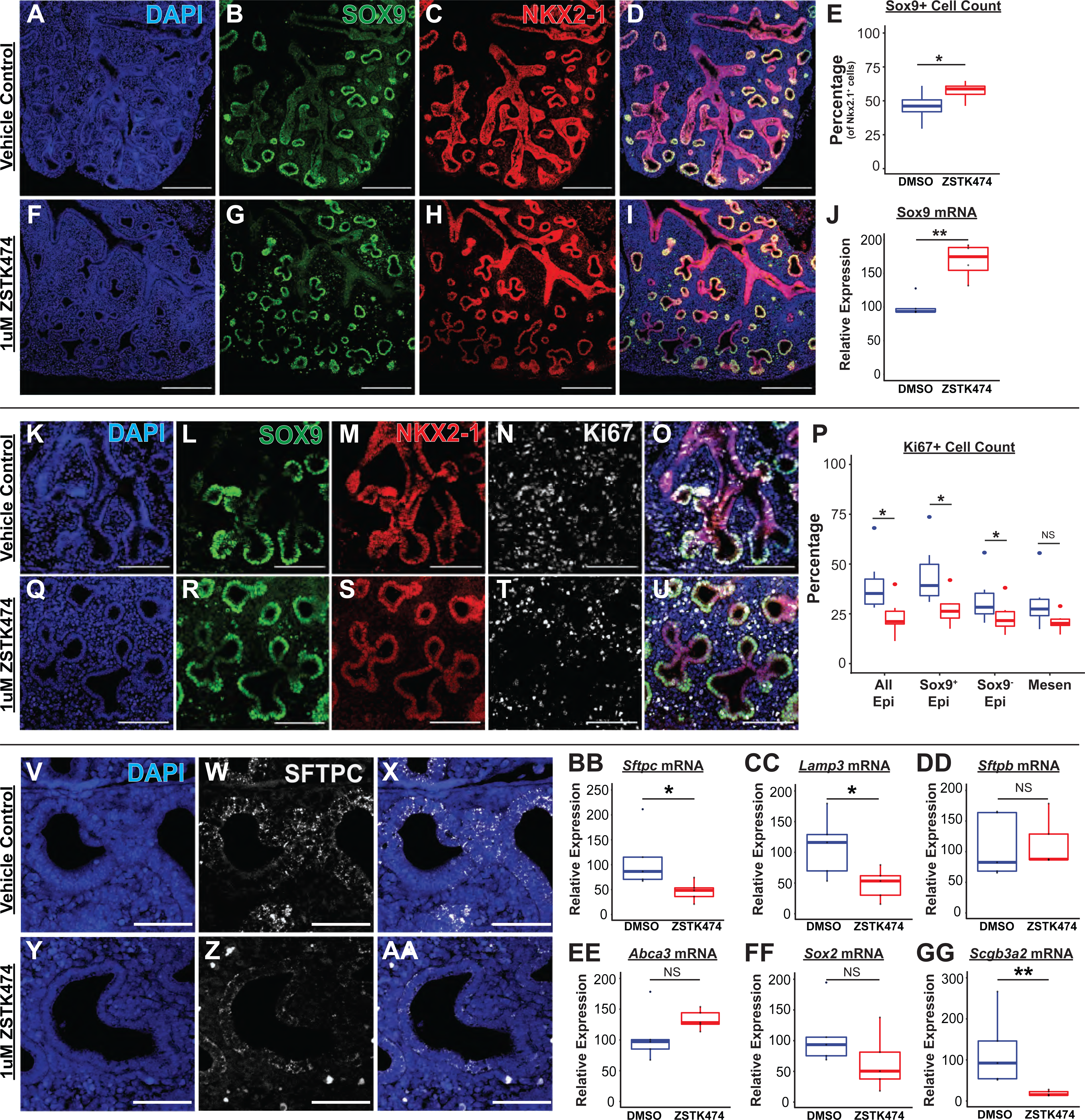
PI3K inhbition increases the number of SOX9+ lung epithelial cells and impairs distal epithelial differentiation in lung explants. (A-I) SOX9+ epithelial cells increase as a proportion of the total NKX2.1+ cell population following PI3K inhibition. (J) Increased *Sox9* mRNA was detected by qPCR. (K-P) The number of Ki67+ proliferating cells was decreased in both the epithelium and mesenchyme following PI3K inhibition. (V-AA) Decreased mature SFTPC protein was detected by immunofluorescence microscopy in distal cells. (BB-CC) Decreased expression of surfactant-associated genes *Sftpc* and *Lamp3* mRNA was detected by qPCR. Scale bars: 200 μm (A-I), 100 μm (K – U). * p < 0.05, ** p < 0.01.

### Epithelial-specific deletion of the *Pi3kca* gene *in vivo* causes in expansion of SOX9+ EPCs and inhibition of epithelial differentiation

To test the hypothesis that epithelial activation of PI3K signaling is required for normal differentiation and proximal-distal patterning of the lung epithelium, we used *Shh*-Cre x *Pi3kca*^f/f^ mice to conditionally delete the *Pi3kca* gene from the embryonic mouse lung epithelium (Harfe et al., 2004; Zhao et al., 2006). Lungs from conditional knockout mice (hereafter referred to as *Pik3ca^Shh^*^Cre^) were harvested at E18.5 and compared to littermate controls (including mice both with and without the Cre allele). The lungs of *Pik3ca^Shh^*^Cre^ mice were remarkably mispatterned, with abnormal cysts lined by NKX2.1+ epithelial cells scattered throughout the lungs (Figure 8A-E, H), with the remainder of the lungs filled with simplified alveolar structures lined by increased numbers of SOX9+ lung epithelial cells (Figure 8C-G, I-P). Perhaps the most notable feature, however, was the marked reduction mature airway epithelium. The number of mature club (SCGB1A1+) or cilated (TUB1A1+) cells present in the lungs of *Pik3ca^Shh^*^Cre^ embyros was significantly reduced, accompanied by significant reduction in the expression of transcripts associated with secretory and ciliated cells, including *Scgb1a1*, *Scgb3a2*, and *Foxj1*. (Figure 8Q-DD and Supplemental Figure 6).

**Figure 8.**
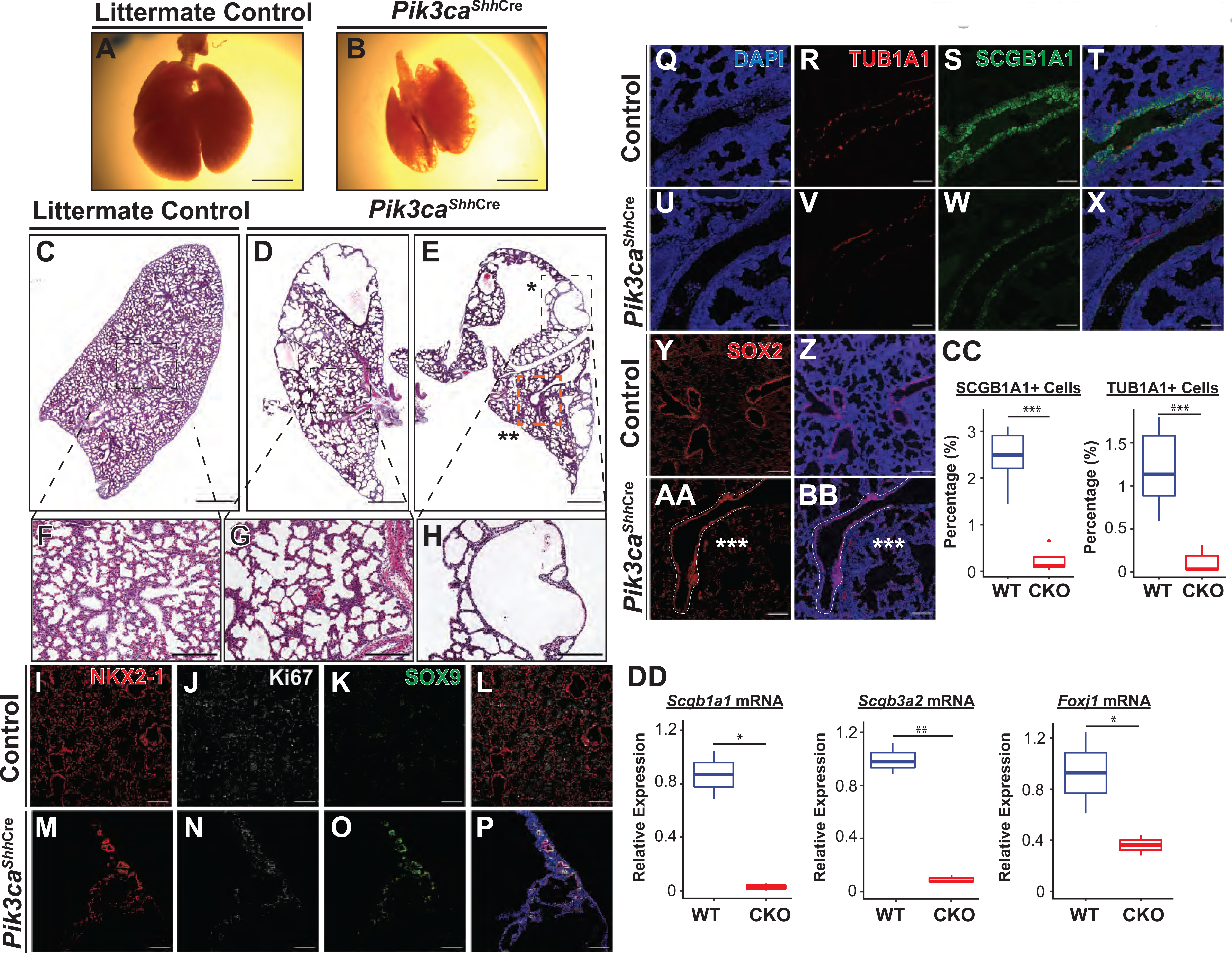
Conditional deletion of *Pi3kca* from the developing lung epithelium causes cystic malformations, persistence of SOX9+ progenitor cells, and impaired proximal lung epithelial development. (A-B) Whole-mount images show cystic areas throughout the lungs of *Pik3ca^Shh^*^Cre^ embryos at E18.5. The lungs are smaller in size compared to littermate controls. (C-H) Widefield imaging of H&E stains shows numerous cystic areas in *Pik3ca^Shh^*^Cre^ and a paucity of conducting airways. The majority of the lung tissue consists of simplified alveolar structures. (I-P) Persistence of the SOX9+ epithelial progenitor cells at E18.5 was observed in *Pik3ca^Shh^*^Cre^, in contrast to littermate controls where few SOX9+ progenitor cells remain. (Q-X, CC) A marked reduction in the number of SCGB1A1+ secretory cells and TUB1A1+ ciliated cells were observed in *Pik3ca^Shh^*^Cre^. The staining intensity of both proteins were also decreased. (Y-BB) The conducting airway epithelium present exhibited SOX2+ staining intensity comparable to wild-type airways. (DD) Decreased expression of transcripts characteristic of secretory and ciliated cells, such as *Scgb1a1*, *Scgb3a2*, and *Foxj1*, were decreased. Scale bars: 2.5 mm (A-B), 500 μm (C-E), 200 μm (F-H), 100 μm (I-P), 50 μm (Q-X). * p < 0.05, ** p < 0.01.

Concordant with our finding in lung explants, these data demonstrate that epithelial-specific PI3K signaling is required for proper proximal/distal patterning of the lung during development. Both *in vivo* and *in vitro* data demonstrate that loss of PI3K signaling causes persistence of the SOX9+ lung epithelial progenitors into late embryonic life, inhibiting normal proximal lung epithelial differentiation and resulting in mispatterning of both the airway and alveolar structures of the lung. Finally, this requirement for PI3K signaling demonstrates that careful comparative analysis using both chromatin accessibility and gene expression data provides higher sensitivity to identify novel regulators of development which may be unrecognized from expression data alone.

## Discussion

The mature mammalian lung is composed of dozens of specialized cell types that are necessary to permit gas exchange and facilitate terrestrial life (Basil et al., 2020; Basil & Morrisey, 2020; Morrisey & Hogan, 2010; Swarr & Morrisey, 2015). While many of the major transcription factors and signaling pathways that drive differentiation of lung cells during development have been defined, mechanisms by which these signals are integrated into the chromatin state to establish and maintain cellular identity remains incompletely understood. Here, we have generated a map of the chromatin accessibility landscape during SOX9+ lung epithelial progenitor cell development, which we have made readily available to the scientific community through the LGEA Web Portal (https://research.cchmc.org/pbge/lunggens/mainportal.html), a product of the NIH LungMap Project (Ardini-Poleske et al., 2017; Du, Guo, Whitsett, & Xu, 2015; Du et al., 2017). Accessible chromatin regions identified in our study are highly correlated with histone post-translational modificiations normally associated with active, functional cis-regulatory elements, and changes in the accessibility of these regions of chromain are associated with changes in developmental gene expression. Changes in chromatin accessibility were preferentially located at putative enhancers, highlighting the importance of targeted chromatin remodeling at select enhancers in the development of the SOX9+ lung epithelial progenitors.

Our combinatorial approach to evaluation of both mRNA and chromatin accessibility data led to the observation that PI3K was a key regulator of EPC development. Accessible regions of chromation, particularly those regions that became more accessible as development proceeded, were significantly enriched nearest genes associated associated with PI3K signaling. Signaling through the PI3K pathway, as measured by pAKT immunostaining was highest in the proximal lung epithelium throughout development. Treatment of embryonic lung explants with PI3K inhibitors increased branching, associated with in increased proportion of SOX9+ lung epithelial cells, *Sox9* mRNA expression, and decreased expression of mature AT2 markers*. In vivo*, a paucity of airways were observed in the *Pi3kca^Shh^*^Cre^ lungs; although the airways present were lined by SOX2+ epithelium, mature secretory and ciliated cell markers were decreased. *Pi3kca^Shh^*^Cre^ mice also demonstrated extensive distal cysts and simplified alveoli, somewhat different than the increased branching observed in lung explants. Both results emphasize the need for an intact gradient of PI3K signaling during development for proper differentiation of the proximal and distal portions of the lung. Although inhibition of PI3K signaling has been previously shown to increase lung branching (at least in some developmental contexts), our results highlight the need for a gradient of PI3K signaling during development for proper differentiation of the proximal and distal portions of the lung (Carter et al., 2014; Metzger, Stahlman, & Shannon, 2008; Metzger, Xu, & Shannon, 2007). Follow-up studies using inducible Cre drivers to inactivate the PI3K pathway at specific stages of development, and examination of developmental time-series following pathway inactivation will be needed to understand the time- and context-dependency of PI3K signaling in lung epithelial development.

Interestingly, expansion of the SOX9+ progenitor cell population was seen in lung explants despite an overall decrease in the number of proliferating (Ki67+) cells in the lung epithelium and mesenchyme. PI3K signaling plays a well-defined role in regulating cell cycle progression, the observed effects were consistent using two PI3K inhibitors, and no toxicities were observed during the 48 hour culture period (Chang et al., 2003). Present observations suggest that inhibition of PI3K signaling restains differentiation of SOX9+ progenitors into more mature cell types, rather than simply increasing the numbers of SOX9+ progenitors through proliferation.

The markedly abnormal lung epithelium seen in *Pi3kca^Shh^*^Cre^ animals imply a critical role in lung development, raising several future questions. First, the receptor/ligand pairs modulating PI3K signaling in the developing lung remain to be defined. FGF signaling, particularly FGF7 and FGF10 signaling through FGFR1/2 has been shown to play a key role in initial outgrowth of the lung bud and subsequent branching morphogenesis. However, abrogation of FGF7 or FGF10 signaling through FGFR1/2 causes apoptosis and failure of lung epithelial outgrowth, not expansion of SOX9+ progenitor cells as observed presently, and thus the phenotypes observed in *Pi3kca^Shh^*^Cre^ animals cannot simply be explained by a downstream effect of the FGF7/10-FGFR1/2 signaling axis; EGF, other FGFs, and IGF are all potential regulators to be explored. Nortably, these result expand on prior studies that have also reported an increae

Second, our data suggest that the PI3K pathway may regulate chromatin state during lung development. Studies from cancer, where aberrant activation of PI3K signaling is common, have shown that pAKT can directly interact with a number of key chromatin regulatory (CR) complexes to direct modulate chromatin state (Yang, Jiang, & Hou, 2019). We speculate that the PI3K signaling pathway serves as the central “switch-board” integrating signals received from diverse receptor ligand-pairs during lung epithelial development, and in turn directly modulates chromatin state, including chromatin accessibility, through interactions between pAKT and key CR complexes to direct lung epithelial differentiation (Figure 9). Future studies investigating the interface of PI3K signaling within the lung epithelium on chromatin state, cellular differentiation, and cell-type identity will be of significant interest in defining these mechanisms.

**Figure 9.**
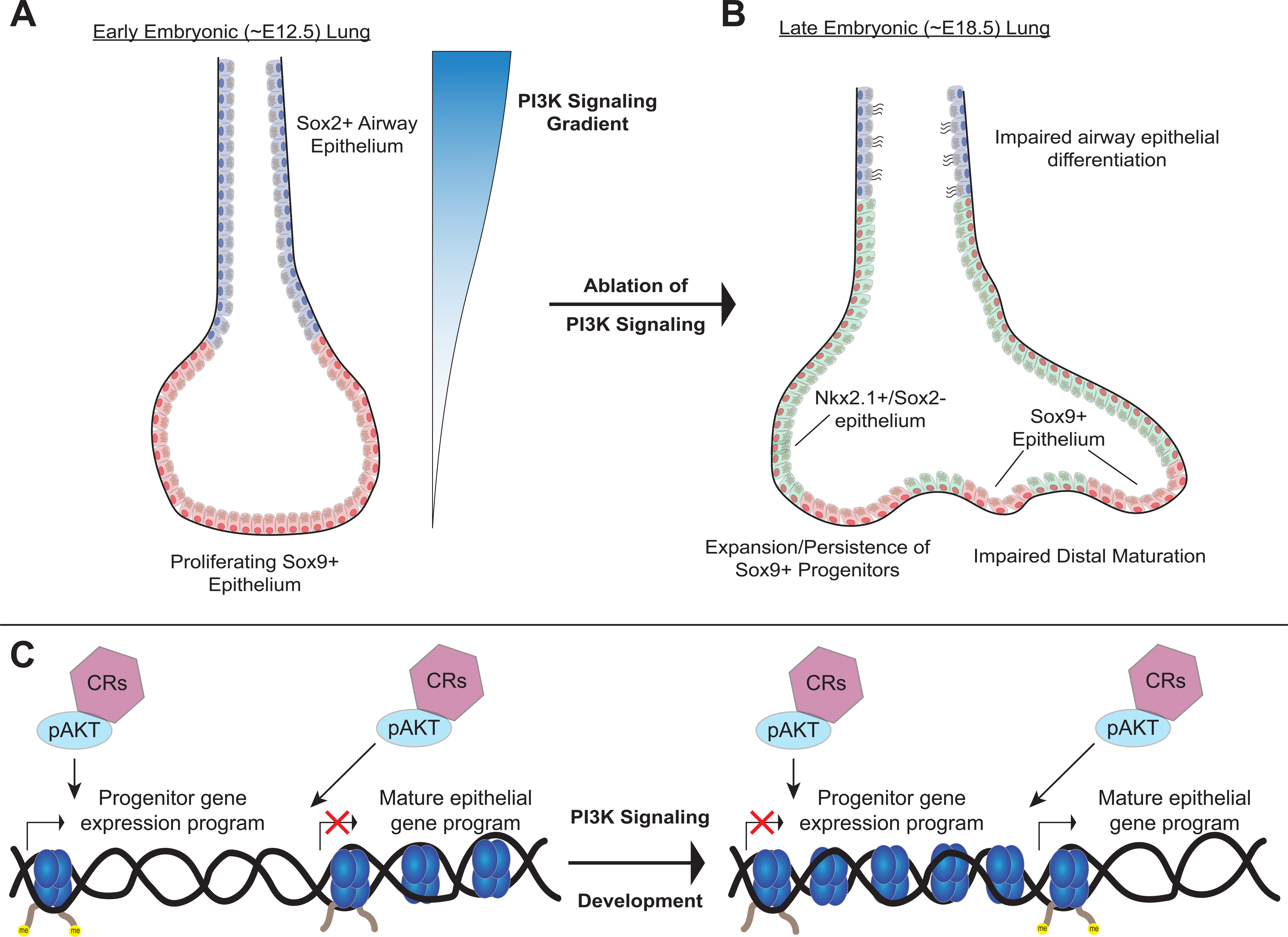
A model of the role of PI3K signaling in the developing lung epithelium. (A) During normal lung epithelial development, a proximal-to-distal gradient of PI3K signaling patterns the lung epithelium, with highest levels in developing conducting airways. (B) Inhibition of PI3K signaling causes expansion or persistence of the SOX9+ progenitor cells, and impairs epithelial differentiation of alveolar and conducting airway epithelial cells. (C) PI3K signaling may directly modulate chromatin accessibility to promote differentiation and pattern cell-type identity during lung epithelial development.

Finally, our data highlight the dearth of data regarding the molecular mechanisms by which chromatin accessibility changes are regulated during differentiation of the SOX9+ EPC and other lung progenitor populations. It remains unknown which of the dozens of proteins and complexes that are known to remodel chromatin are specifically required for development and differentiation of the lung epithelium at one specific timepoint, let alone globally. Studies in other tissues imply that specific CR complexes interface with major developmental transcription factors to mediate chromatin accessibility, and that chromatin accessibility changes are an integral part of cell-type identity. Given the promising progress of epigenomic therapies in cancer (Bates, 2020), there is an urgent need for *in vivo* genetic studies targeting specific CR complexes and CRISPR-based epigenome engineering strategies (Holtzman & Gersbach, 2018) to define the epigenomic mechanisms of lung homeostasis and disease.

## Methods

### Animal Husbandry

Experiments were performed on a mixed C57BL/6, CD-1 background. All animal work was performed under the approval and guidance of the Cincinnati Children’s Hospital Institutional Animal Care and Use Committee (IACUC). The Tg(Sox9-EGFP)^EB209Gsat^ mouse line (MGI ID: 3844824) was used for isolation of SOX9+ epithelial cells, using the methods described below (Gong et al., 2003). The *Shh*^tm1(EGFP/cre)Cjt)^ (JAX stock # 055622) and *Pik3ca*^tm1Jjz^ (JAX stock #017704) lines were used to conditionally delete Pik3ca from the developing lung epithelium (Harfe et al., 2004; Zhao et al., 2006).

### Single-Cell RNA Sequencing Analysis

Publicly available scRNA-seq data from EpCAM sorted e11 mouse lungs were combined with publicly available e16 whole lung scRNA using Seurat’s *FindIntegrationAnchors*, using only genes found in both datasets (Kuwahara et al., 2020) After integration, SeuratWrappers and Monocle 3 were used to create a linage trajectory after indicating E11 epithelial cells as the root of the model, minimal_branch_len=18. After sub-setting out all epithelial cells, gene modules were calculated using Monocle3’s *find_gene_modules*, resolution=1e-3, with *graph_test* results as input, q_value < 0.01. Unless indicated all default settings for Mononcle3 were used. Marker genes for e16 distal and proximal epithelial cells were generated by comparing only epithelial clusters using Seurat’s *FindMarkers*. Genes in gene modules of interest were analyzed by Toppfunn’s functional enrichment analyses to determine biological process related to specific modules.

### SOX9+ Lung Epithelial Progenitor Cell Isolation

Timed matings were performed by placing a Tg(Sox9-EGFP)^EB209Gsat^ male reporter mouse with a wild-type female, and the presence of a copulation plug the following morning was used to define gestational day (GD) 0.5. Lungs were microdissected from embryos isolated from pregnant females at E11.5 and E16.5 (n = 3 at each timepoint). Lung lobes were isolated from trachea, eosphagus and surrounding tissue, minced with fine scissors, and then were digested with 1mL 10X TrypLE (Gibco) for 5-10 minutes at 37C with agitation. Cells were filtered (70um and 35um filters, respectively) and washed with FACS buffer (HBSS buffer(Gibco), 2% FBS, 25mM HEPES buffer pH 7.0, 2mM EDTA pH 7.4) twice. Next, cells were labeled with anti-CD326 (eBioscience #17-5791-82), anti-CD31 (eBioscience #25-0311-82) and anti-CD45 (eBioscience #15-0451-82) at a 1:200 dilution for 10 minutes at 4C. Cells were then washed again, and treated with 7-AAD (BioLegend) (5uL per 100uL FACS buffer). SOX9+ lung epithelial cells were isolated by sorting for GFP+/CD326+/CD31-/CD45-/7-AAD-using a BD/FACSAria II cell sorter. Cells were sorted directly into Trizol (ThermoFisher Scientific) for RNA isolation, or enriched cell-culture media (HBSS buffer (Gibco), 25mM HEPES buffer, 50% FBS) for ATAC-Seq library preparation.

### RNA-Seq Library Preparation, Sequencing & Bioinformatic Analysis

Isolated SOX9+ lung epithelial cells sorted directly into Trizol were sent to GeneWiz, LLC for RNA isolation and library preparation. After Trizol/chloroform extraction, RNA isolation was completed with the RNeasy Micro Plus kit per manufacturer’s instructions (Qiagen). SMART-Seq v4 Ultra-Low Input RNA kits (TaKaRa Bio) were used to generated polyA selected libraries from total RNA. Sequencing was performed on an Illumina HiSeq platform, using 2x150bp configuration. Quality trimming and adapter clipping of sequencing reads was performed with Trimmmatic (Bolger, Lohse, & Usadel, 2014). Read quality was assessed before and after trimming using the FastQC program. Reads were then aligned against the mouse reference genome (mm10) using the STAR aligner (Dobin et al., 2013). Duplicate reads were flagged and removed using the MarkDuplicates program from Picard tools. Per-gene read counts for GENECODE release M12 (GRCm38.p5) were computed using the Rsubread R package, with duplicate reads removed (Frankish et al., 2019; Liao, Smyth, & Shi, 2019). Gene counts represented as counts per million (CPM) were first nominalized using the TMMmethod in the edgeR R package, and genes with 25% of samples with a CPM<1 were removed. The data were transformed using the VOOM function from the limma R package (Law, Chen, Shi, & Smyth, 2014). Differential gene expression was performed using a linear model with the limma package. Heat maps and principal component analysis plots were generated in R. GO enrichment analysis for the biological process was performed using the R package GAGE (Luo, Friedman, Shedden, Hankenson, & Woolf, 2009). Expression of select gene targets were displayed using the boxplot function of ggplot2() in R with tpm counts generated by Stringtie. (Pertea et al., 2015; Wickham, 2016)

### ATAC-Seq Library Preparation, Sequencing & Bioinformatic Analysis

Isolated SOX9+ lung epithelial cells were sorted into HBSS buffer (Gibco) containing 50% FBS and 25mM HEPES buffer (Gibco). Nuclei were isolated, the transposase reaction was performed with Nextera Tn5 transposase (Illumina), and PCR amplification was performed according to previously published protocols (Buenrostro, Wu, Chang, & Greenleaf, 2015). Sequencing was performed by GeneWiz, LLC on an Illumina HiSeq platform, using 2x150bp configuration. Reads were quality filtered and adapters were trimmed using Trimmomatic, version 0.39 (Bolger et al., 2014). Reads were aligned to mouse genome mm10 using bowtie2 (Langmead & Salzberg, 2012). Duplicated reads were removed with picard tools, and blacklisted regions were filtered out using bedtools (Amemiya, Kundaje, & Boyle, 2019). Peak calling was performed with MACS2 (version 2.2.1.20160309) using the additional parameters “—nomodel”, “—shift -100”, and “—extsize 200”. The MACS2 “bdgdiff” function was used to identify differentially accessible regions. A q-value (FDR) of 0.05 was used to determine significance for both analyses. Peak annotation, gene ontology analysis of nearest genes, and quantification of ENCODE ChIP-Seq read density by ATAC-seq peak location analyses were performed using Homer (Heinz et al., 2010). The R software package was used to measure and display relationships between ATAC-Seq peak location and nearest gene neighbor expression values. ChIP-Seq data for H3K4me1, H3K4me3, and H3K27ac were accessed either directly from the ENCODE Project website (www.encodeproject.org) or via the UCSC genome browser (https://genome.ucsc.edu/) (Consortium, 2012; Davis et al., 2018; Sloan et al., 2016). A list of ENCODE accession numbers and generating laboratories for these datasets appears in Supplemental Table 7. Genomic interval plots were generated using the UCSC genome browser, using the GRCm38/mm10 genome builds. Paired expression and chromatin accessibility modeling was performed using the methods and software described previously (Duren et al., 2017). Stringtie was used to generate tpm based expression matrices as imput into PECA from merged bam files for all replicates at each developmental timepoint (Pertea et al., 2015). Replicated ATAC-Seq peaks were used as the chromatin accessibility input to PECA.

### Lung Explant Culture Model

Embryonic lungs were harvested by micro-dissection at E12.5 following timed-matings between CD1 wild-type mice. The isolated whole lung explants were placed on Millicell Cell Culture Inserts (1.0uM PET, Millipore, Ref# MCRP06H48) suspended on 700uL of DMEM/F12K culture medium (Gibco, Ref# 12634-010) supplemented with 10% FBS and 1% Anti-anti (Gibco, Ref# 15240-062) and placed in a 37C incubator (5% CO2). The lung explants were treated with one of two different pan-class I PI3K pathway inhibitors, ZSTK474 (Cell Signaling Ref# 13213) at concentration of 1uM, or BYL719 (Selleckchem Cat# S2814) at concentration of 1uM, 5uM or 10uM for 48hrs. Controls were treated with 0.1% DMSO. Whole-mount images were acquired at 0, 24 and 48 hours in culture using a Leica MZ16 FA fluorescent stereomicroscope and DFC7000T camera system. Branching was quantified by counting the number of terminal branches visible and calculating the percentage increase in number of terminal branches in the inhibitor-treated explants compared to the controls. Each lung explant was either then fixed, emedded and used for histological studies as described below, or was used to generate a single-cell suspension for RNA analysis by qPCR (see below).

### Histology & Immunofluorescence Microscopy

Tissues were fixed in 4% paraformaldehyde, embedded in paraffin wax, and sectioned at 5um intervals. Slides were stained with hematoxylin using standard protocols, as previously described (Herriges et al., 2014; Swarr et al., 2019). Immunofluorescence was performed using the following antibodies: anti-NKX2.1 (guinea pig, Seven Hills, 1:500), anti-SOX9 (rabbit, Millipore, 1:100), anti-KI67 (mouse, BD Pharmigen, 1:100), anti-SFTPC (goat, Santa Cruz, 1:200), anti-SCGB1A1 (rabbit, Seven Hills, 1:500), anti-TUB1A1 (mouse, Sigma Aldrich, 1:1000), and anti-SOX2 (mouse, Santa Cruz, 1:100). Immunofluorescence for phosphorylated-ATK (pAKT) was performed with anti-pAKT (rabbit, Cell Signaling, 1:100), followed by multi-step detection with the Biotin-Tyramide signal amplification kit (Perkin Elmer) with a 15 minute exposure time. TUNEL staining was performed using the TACS-XL Blue Label kit, per manufacturer’s instructions (Trevigen). All slides were mounted with Prolong Gold Antifade medium (Invitrogen), and were then imaged on either a Nikon Eclipse 90i widefield, or Nikon A1R GaAsP Inverted Confocal Microscope. Cell type quanitification based on immunofluorescence microscopy images was performed using the Nikon Elements Advanced Analysis software suite.

### Whole-mount Imaging

Tracheal lung tissue isolated at E12.5, E13.5 and E14.5 was subject to whole-mount immunofluorescence staining as described (Sinner et al., 2019). Embryonic tissue was fixed in 4% PFA overnight and then stored in 100% MeOH at -20^0^C. Whole-mounts were permeabilized in Dent’s Bleach (4:1:1 MeOH: DMSO: 30%H2O2) for 2 hours, then taken from 100% MeOH to 100% PBS. Following washes, tissue was blocked in a 5% (w/v) blocking solution for two hours and then incubated, overnight, at 4^0^C in primary antibody solution. After washes in PBS, whole-mounts were incubated with a secondary antibody overnight at 4^0^C. Samples were then washed, dehydrated and cleared in Murray’s Clear. Images of whole-mounts were obtained using confocal microscopy (Nikon A1R). Imaris imaging software was used to convert z-stack image slices obtained using confocal microscopy to 3D renderings of whole-mount samples.

### Quantitative real-time PCR

A single-cell suspension was generated, using the methods described above, from either lung explant cultures or embryonic mouse lungs isolated at E18.5 from Pik3ca^ShhCre^ embryos and littermate controls. Epithelial cells were isolated using sheep anti-rat Dynabeads (Thermo Fisher) with rat anti-mouse CD326 antibody (Thermo Fisher). Total RNA was subsequently isolated from the bead-sorted epithelial cells using Trizol (Invitrogen), per the manufacturer’s protocol. cDNA was synthesized from total RNA by using the SuperScript IV strand synthesis system (Invitrogen). Quantitative real-time PCR was performed using the SYBR Green system (Applied Biosystems) with primers listed in Supplemental Table 8. *Gapdh* or *Tbp* expression values were used to control for RNA quantity. Two-tailed Student’s T-Test was used for all comparisons involving two groups.

## Acknowledgements

We would like to acknowledge the assistance of the Research Flow Cytometry Core and Confocal Imaging Core at Cincinnati Children’s Hospital Medical Center. We also would like to thank Jeffrey Whittset, MD for his careful review of this manuscript, and Lauren Leesman for assistance in performing the whole-mount imaging presented here. We are thankful for the generous grant support from the following sources: National Institutes of Health grants K08HL130666 (DTS), K08HL140178 (WJZ), R01HL144774 (D. Sinner), and CCHMC Parker B. Francis Fellowship Award (DTS).

## Author Contributions

D.K., S.F., M. Gillen, and D. Sinner, D.T.S all performed experiments related to this manuscript. D.K, S.F., J.S, M. Guo, and D.S. performed the data analysis in this manuscript. D.K., S.F., W.J.Z, D. Sinner, J.S., and D.T.S. all contributed to writing the manuscript.

## Additional Information

### Data Access

The data sets from this study have been submitted to the NCBI Gene Expression Omnibus (GEO; https://www.ncbi.nlm.nih.gov/geo/) under accession number: PENDING.

## Supplemental Materials

**Supplemental Figure 1.**
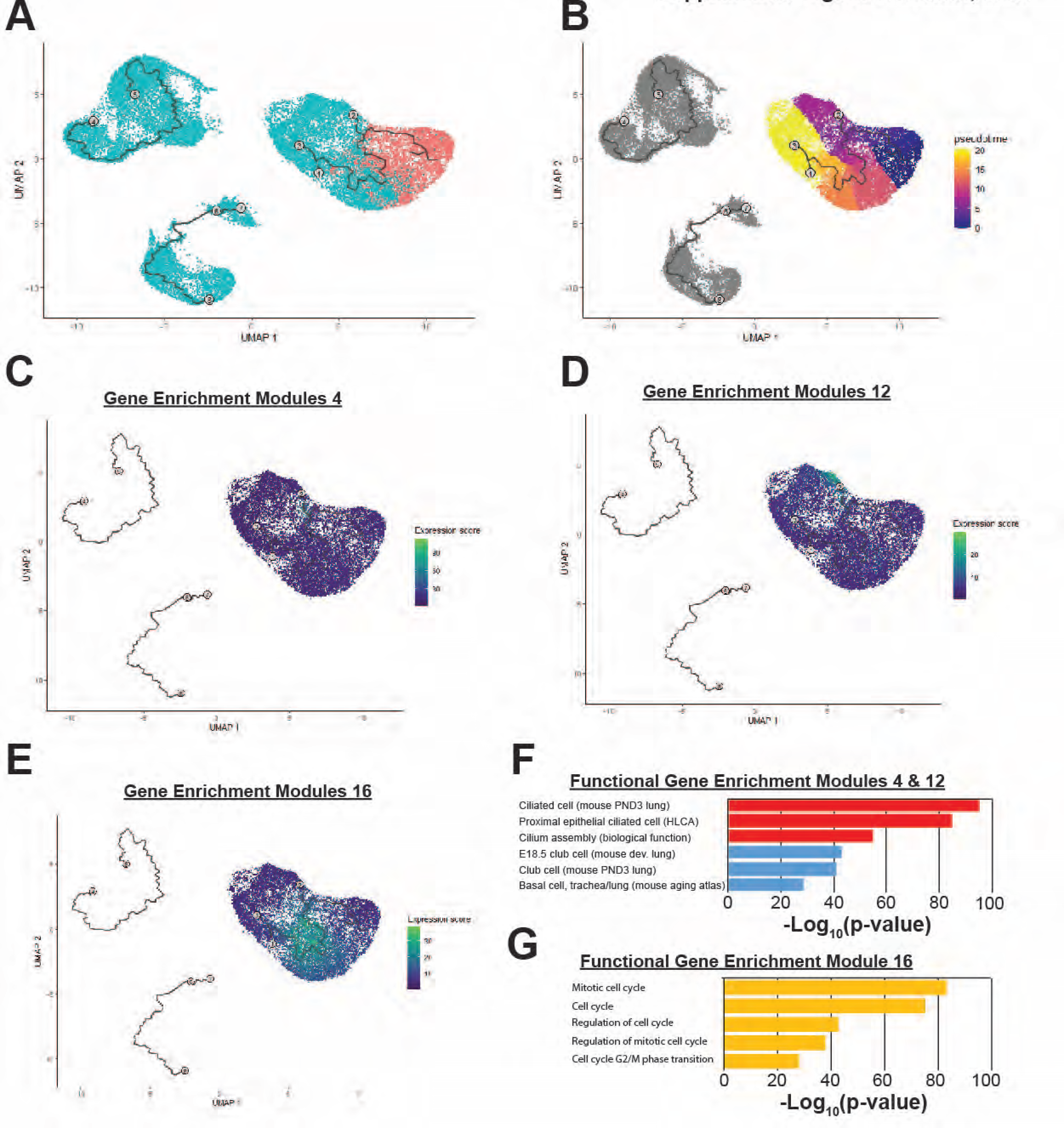
Single-Cell Analysis of Mouse Embryonic Lung. (A) Sample batches are indicated (red – E11.5 lung epithelium blue – E16.5 lung epithelium). (B) Monocle pseudotime heatmap is shown. (C-D) Gene enrichment modules 4 and 12 are highly enriched for components of the ciliated and secretory cell epithelium respectively. (E) Gene enrichment module 16 are highly enriched in genes involved in cell cycle regulation, and cells expressing these targets are found in greatest numbers at the center of the pseudotime pathway. (F-G) Gene set enrichment categories for modules 4 & 12 (F) and module 16 (G) are shown.

**Supplemental Figure 2.**
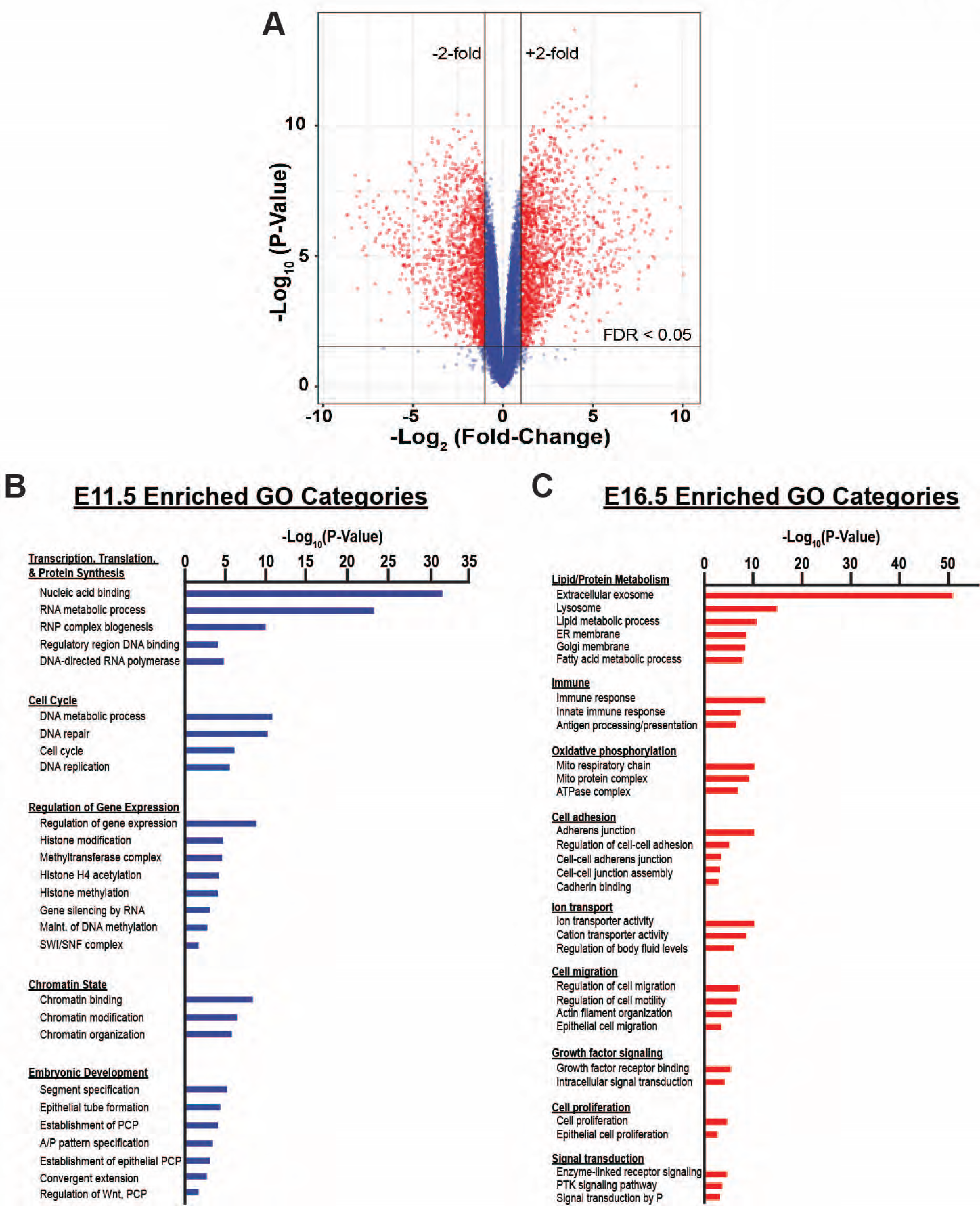
Bulk RNA-Seq Analysis of the developing SOX9+ EPC population. (A) A Volcano plot for genes differentially expressed between E11.5 and E16.5 in SOX9+ EPCs is shown. (B-C) Gene ontology analysis demonstrates that differentially expressed genes at E11.5 are enriched in functions involved with cell proliferation, protein synthesis, and “signaling pathways regulating pluripotency” (B), whereas genes with increased expression levels at E16.5 are enriched in functions associated with the mature distal lung epithelium, such as surfactant production (e.g. “phagosome”, “lysosome”, “sphingolipid metabolism”, “protein export), epithelial barrier function (e.g. “ECM-receptor interaction”, “focal adhesion”, “collective duct secretion”), and a switch in energy metabolism (e.g. “oxidative phosphorylation”, “citrate/TCA cycle”) (C).

**Supplemental Figure 3.**
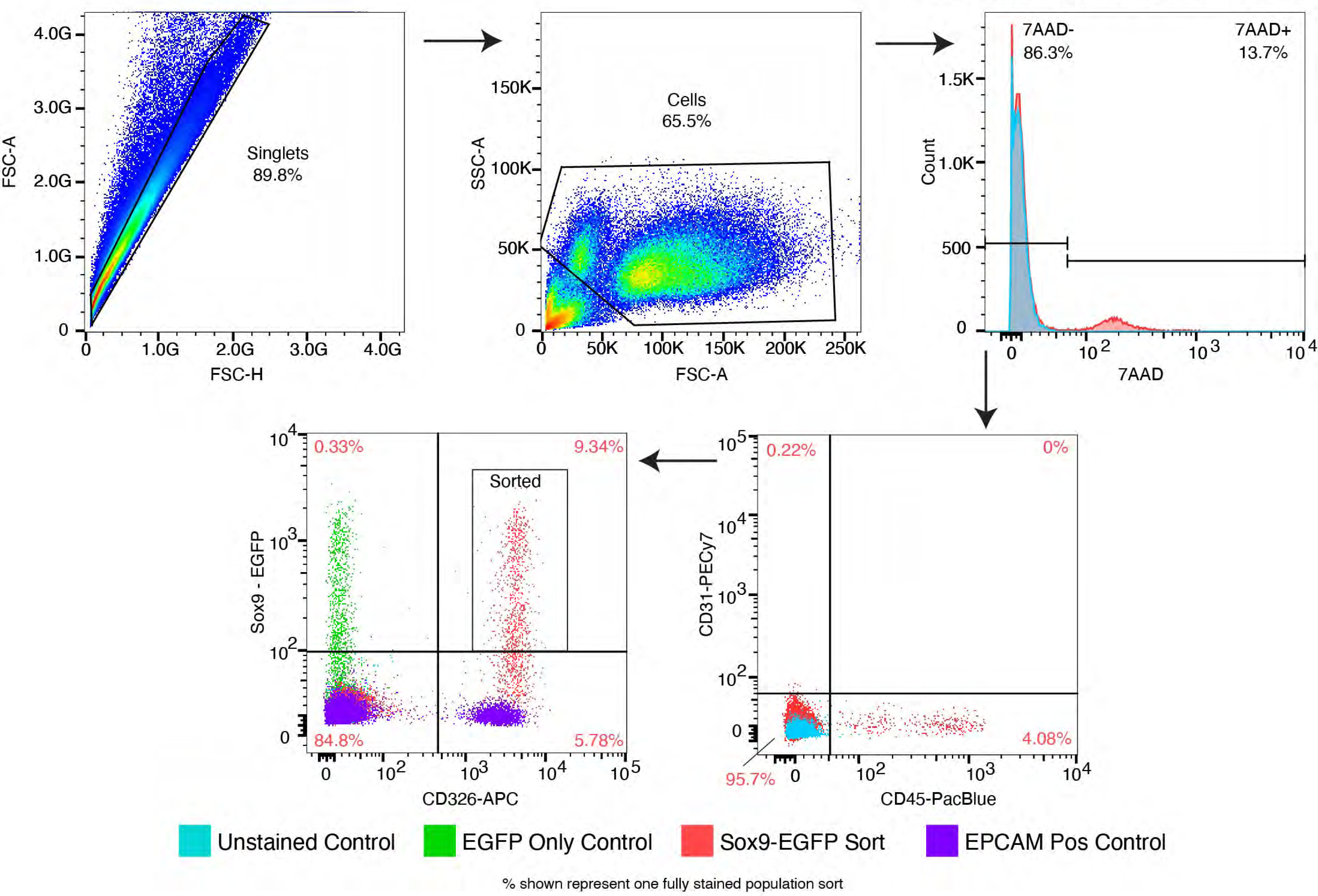
Isolation of SOX9+ EPCs by Fluorescence-Activated Cell Sorting (FACS). SOX9+ EPCs were isolated from a single-cell suspension of whole lung at E11.5 and E16.5 from EGFP-SOX9 reporter mice by sorting for GFP^+^, CD326^+^, CD45^-^, CD31^-^, 7-AAD^-^, to positively select for SOX9+ epithelial cells and negatively select for immune, endothelial, and dead cells, respectively.

**Supplemental Figure 4.**
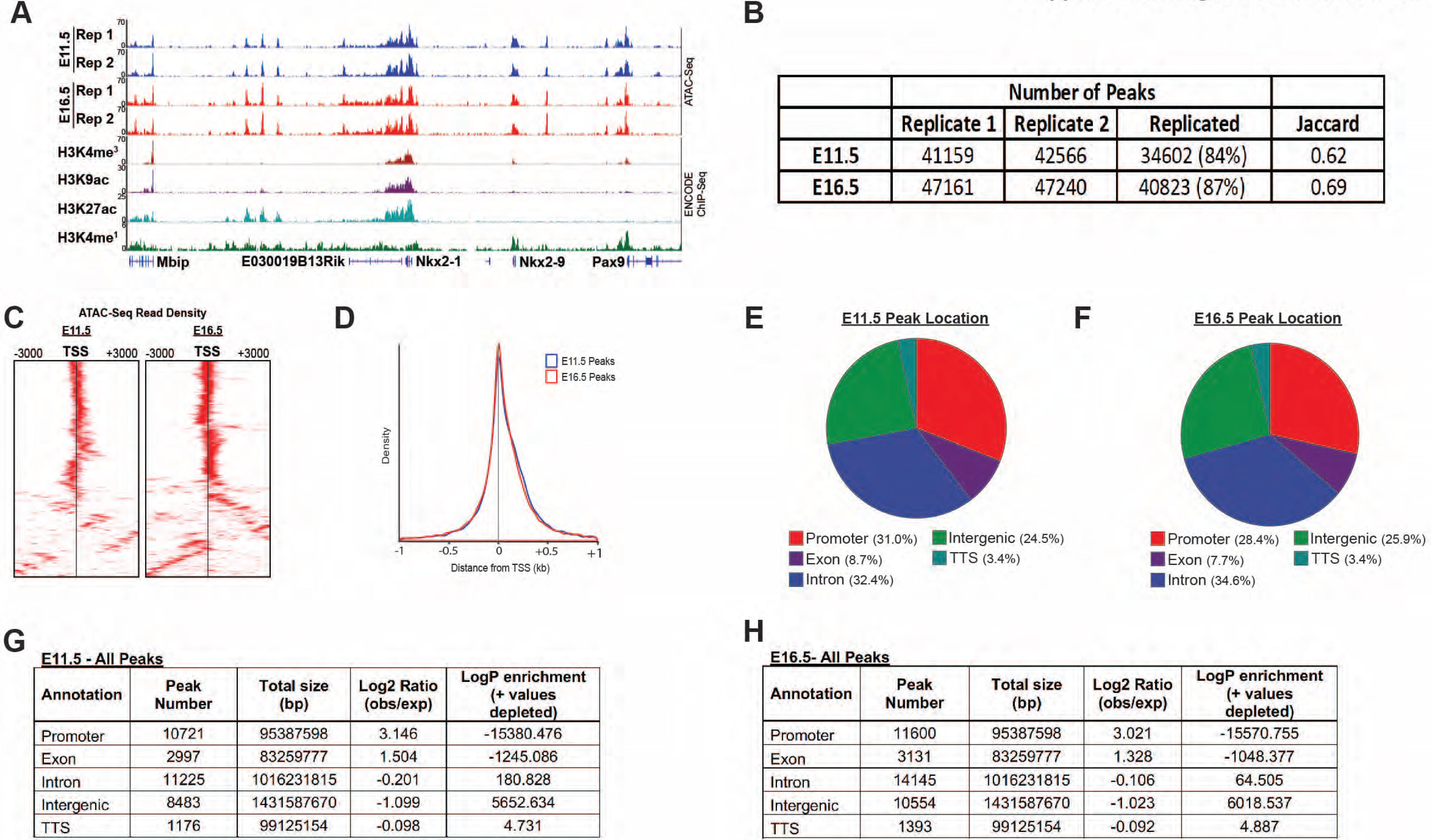
ATAC-Seq Peak Characteristics. (A) A representative locus for a well-expressed gene (*Nkx2.1*) shows the high degree of consistency between biological replicates, and strong correlation between open regions of chromatin with histone post-translational modifications correlated with active cis-regulatory regions. (B) The number of peaks identified by MACS2 for each biological replicate, and replication statistics, are shown. (C) Heatmaps display the high-degree of enrichment for ATAC-seq reads within and immediately surrounding transcriptional start-sites (TSSs). (D) ATAC-req density is plot as a function of distance from the transcriptional start site (TSS). (E-H) The distribution of ATAC-Seq peaks by genomic location is shown.

**Supplemental Figure 5.**
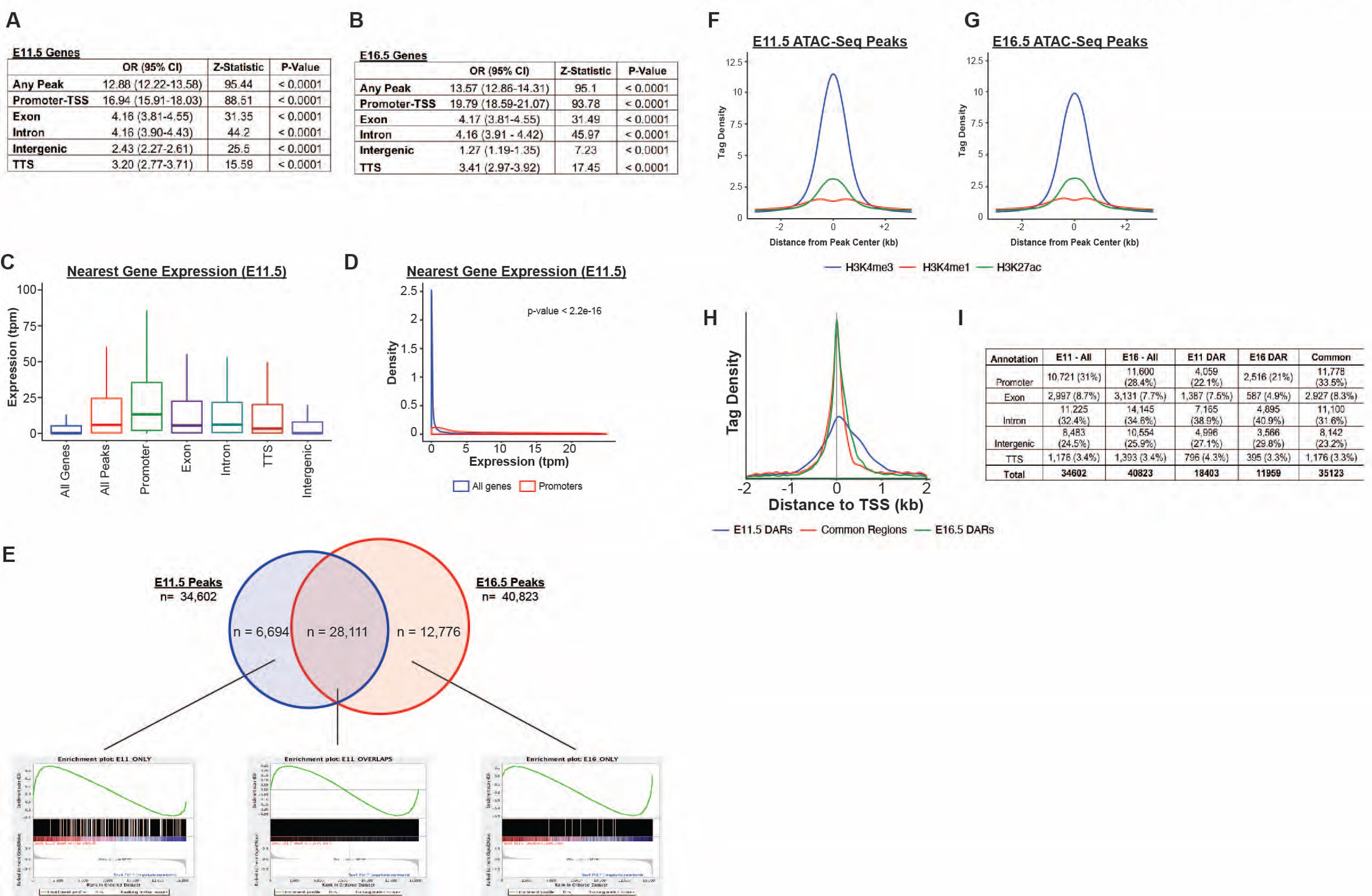
Bulk RNA-Seq Analysis of the developing SOX9+ EPC population. **(A-B)** The relationship between gene expression and the presence of a nearby region of accessible chromatin is displayed. Data is shown as the odds that a gene is expressed, compared to the set of all genes, for those genes identified as the nearest neighbor to a region of accessible chromatin (Any Peak). These relationships are further broken down by the types of genomic region in which the accessible region falls. (C-D) The expression levels for genes nearest to any region of accessible chromatin (All Peaks), or peaks corresponding to a particular genomic region are shown for E11.5, plotted as either a bar-whisker plot (C) or density plot (D) (data for E16.5 is shown in main Figure 3E-F). (E) Gene-set enrichment analysis for peaks unique to E11.5 or peaks unique to E16.5 reveals an increased number of genes with higher levels of expression at that same developmental timepoint. In contrast, peaks identified at both developmental timepoints do not show a consistent direction of differential expression. (F-G) Density plots for H3K4me^3^, H3K4me^1^, and H3K27ac ChIP-Seq read density are displayed relative to the center of identified ATAC-Seq peaks at E11.5 (F) and E16.5 (G). (H) Density of ATAC-Seq reads associated with E11.5 DARs, E16.5 DARs, and common regions relative to transcriptional start sites (TSSs) are shown. (I) The distribution of E11.5 DARs, E16.5 DARs, and common regions by genomic location is shown.

**Supplemental Figure 6.**
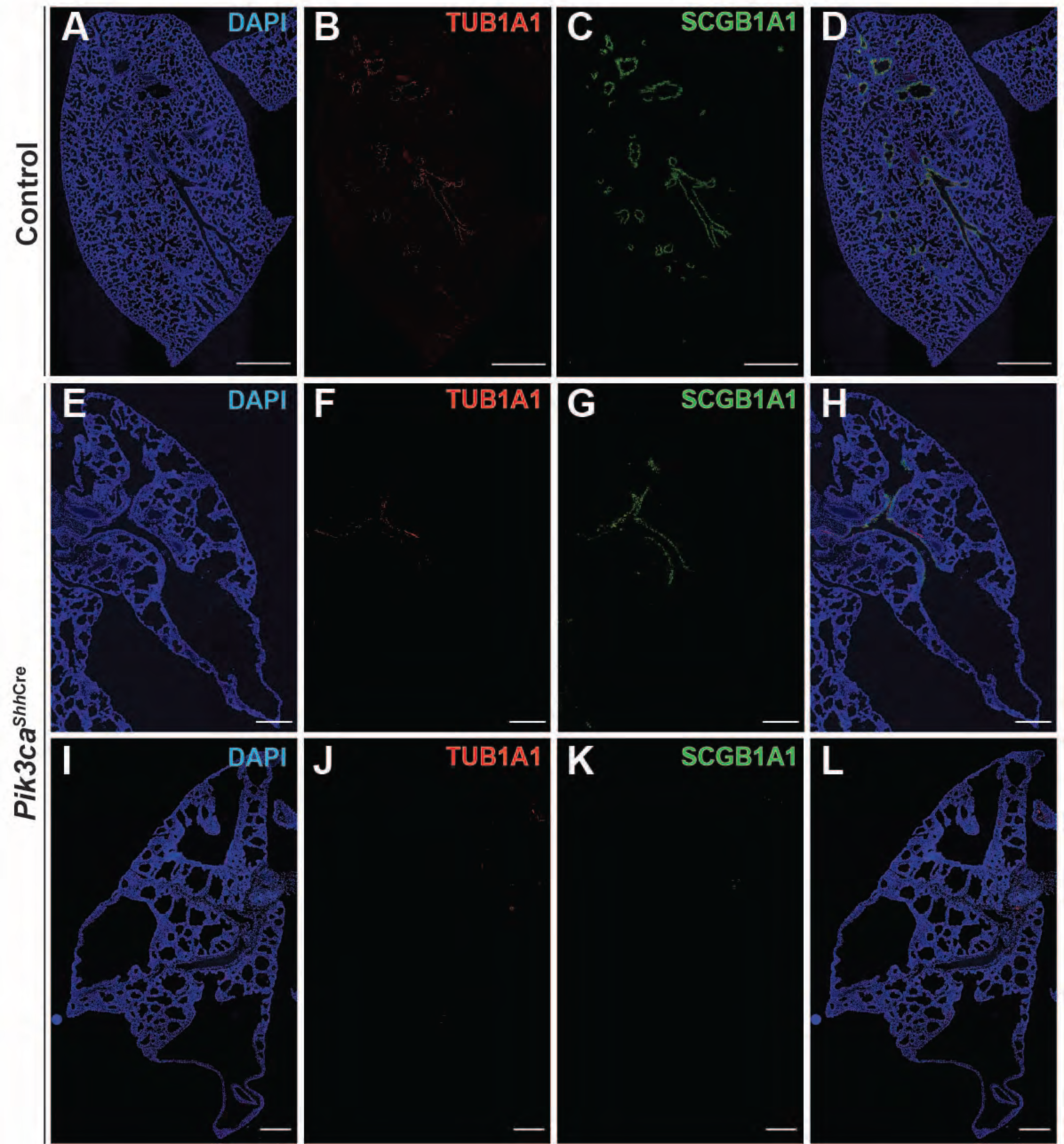
Epithelial-specific loss of *Pik3ca* is associated with a marked reduction of conducting airway epithelium in the lung. (A-D) Lungs isolated from wild-type littermate controls reveal numerous conducting airways throughout the lungs, lined by ciliated (TUB1A1+) and secretory (SCGB1A1) cells. (E-L) In contrast, in lungs isolated from *Pik3ca^Shh^*^Cre^ embryos, the number of conducting airways was either severely diminished (E-H) or essentially absent from whole-lung sections (I-L). Scale bars, 250um.

## Supplemental Tables

**Supplemental Table 1. Differentially Expressed Genes.** Differential gene expression measured by RNA-Seq for E11.5 versus E16.5 SOX9+ epithelial progenitor cells (EPCs). A positive logFC indicates increased expression at E16.5 (compared to E11.5), and vice versa. All comparisons with adjusted p-value < 0.05 are shown, sorted by adjusted p-value.

**Supplemental Table 2. Annotated E11.5 ATAC-Seq Peaks.** ATAC-Seq peaks for E11.5 SOX9+ EPCs are listed, as identified by MACS2 and annotated using HOMER.

**Supplemental Table 3. Annotated E16.5 ATAC-Seq Peaks.** ATAC-Seq peaks for E16.5 SOX9+ EPCs are listed, as identified by MACS2 and annotated using HOMER.

**Supplemental Table 4. Annotated E11.5 Differentially Accessible Regions.** Regions of chromatin that were identified to have statistically significant increases in chromatin accessibility at E11.5 (compared to E16.5) are listed, with peak annotations provided by HOMER.

**Supplemental Table 5. Annotated E16.5 Differentially Accessible Regions.** Regions of chromatin that were identified to have statistically significant increases in chromatin accessibility at E16.5 (compared to E11.5) are listed, with peak annotations provided by HOMER.

**Supplemental Table 6. Annotated Common Accessible Regions.** Regions of chromatin that were accessible at both timepoints and did not undergo significant changes in chromatin accessibility between the two timepoints are listed, with peak annotations provided by HOMER.

**Supplemental Table 7. ENCODE Data References**. The accession numbers, experimental data, and generating laboratory for the ENCODE datasets used in this manuscript are listed.

**Supplemental Table 8. Real Time (RT)-PCR Primer Sequences**. The oligonucleotide sequences used for RT-PCR primers in this manuscript are listed.

